# IL-38 limits alloreactivity through modulating myeloid and T-cell activation

**DOI:** 10.64898/2026.06.19.728400

**Authors:** Anastasiia Kiprina, Wenqing Xu, Igor Mačinković, Nicola Böffinger, Dmitry Namgaladze, Kristin Jordan, Mohammed A.F. Elewa, Anne-Claire Jacomin, Ivan Kur, Blerina Aliraj, Katharina Imkeller, Bernhard Brüne, Andreas Weigert

## Abstract

Interleukin-38 (IL-38) is a cytokine of the IL-1 cytokine family that promotes the resolution of inflammation. Resolution mechanisms comprise the induction or recovery of immune tolerance that is lacking in various acute and chronic inflammatory pathologies, including Graft-versus-Host Disease (GvHD). The role of IL-38 in the context of immune tolerance, its primary immune cell targets and underlying molecular mechanisms are not defined. In this study, we investigated the impact of IL-38 on human alloreactivity and in a mouse model of acute GvHD. Our data suggests that monocytes differentiating into macrophages are the main cellular target of IL-38. Specifically, IL-38 reduces antigen presentation capacity in differentiating monocytes through an IL-1 family receptor-independent mechanism, which subsequently avoids T-cell activation. In parallel, IL-38 ameliorates inflammation in allogeneic settings in human and murine GvHD models by promoting the expansion of regulatory T-cells. Our findings indicate that IL-38 promotes immune tolerance during alloreactivity by affecting myeloid cells and T-cells.

## Introduction

Acute Graft-versus-Host-Disease (aGvHD) is a severe complication occurring after allogeneic stem cell transplantation (allo-HST)^1^, which can lead to massive organ damage^2^ and result in poor survival of patients^3^. GvHD is mediated by alloreactive donor (the graft) T-cells, which recognize the recipient cells (the host) as foreign. Major histocompatibility complex (MHC)-mismatch between donors and recipients is a crucial factor determining disease and survival^4^. The development of GvHD includes 3 phases^5^: (1) tissue damage after conditioning alongside the release of pro-inflammatory cytokines, damage-associated molecular pattern molecules (DAMPs), and pathogen-associated molecular pattern molecules (PAMPs)^6^, (2) priming, activation, and proliferation of the donor T-cells^7^, and (3) the migration of activated lymphocytes to peripheral organs causing destruction^7^. T-cell activation in this context is facilitated not only by direct recognition of mismatched MHC molecules, but also by antigen-presenting cells (APCs) bearing immunogenic host antigens^8^.

During the last decades, the prophylaxis and treatment of GvHD were notably advanced^9^. Currently, prophylaxis heavily relies on post-transplant cyclophosphamide (PTCy). Application of PTCy has revolutionized outcome in patients receiving haploidentical HST^10,11^, and recently has shown benefit for the prevention of GvHD after mismatched unrelated donor (MMUD) HCT^12^. Current developments in the prevention of acute GvHD are, e.g., application of abatacept with calcineurin inhibitors/methotrexate (CNI/MTX), which demonstrated improved survival in patients receiving MMUD HCT or HLA-matched unrelated donor (MUD) HCT^13^. Unfortunately, PTCy is still associated with acute organ failure^14^ and high incidents of cytomegalovirus (CMV) reactivation^15^, and abatacept has not shown potential in chronic GvHD^16^. Standard treatment of established GvHD is based on systemic corticosteroids^17^. Moreover, Janus kinase (JAK) inhibitors have emerged as an effective therapeutic agent against acute and chronic GvHD^18^. However, despite these prevention and therapeutic strategies, 30% to 60% patients still develop moderate to severe cases of GvHD^9^. Therefore, new treatment or prevention strategies, ideally inducing immune tolerance against host antigens rather than generalized immunosuppression, are needed.

IL-38, a member of the IL-1 cytokine family, is proposed as an IL-1-family receptor antagonist that, similar to other IL-1 family receptor antagonists such as IL-1RA and IL-36RA, limits inflammation in various conditions^19–22^. A specific IL-38 receptor is so far unknown. IL-38 was initially shown to bind to the IL-1 receptor IL-1R1^23^ and the IL-36 receptor IL-1R6^24^. However, recent evidence suggests that IL-38 does not efficiently antagonize these receptors^25^. An alternative receptor might be IL-1RAPL1, which is expressed by macrophages and certain T-cells upon activation^26^. Importantly, not only the main receptor, but also critical cellular and molecular targets of IL-38 are poorly defined. However, IL-38 seems to be uniquely equipped among IL-1 family receptor antagonists to promote the resolution of inflammation^27^. Since full resolution of inflammation requires maintaining and/or reestablishing antigen tolerance, we investigated the potential role of IL-38 in this process, using *in vitro* and *in vivo* GvHD models. Moreover, we sought to identify major cellular targets of IL-38.

## Results

### IL-38 suppresses inflammatory and cytotoxic activity of T cells in mixed lymphocyte reactions

To identify major cellular immune targets of IL-38, we aimed at assessing the impact of IL-38 on leukocyte activation in a complex setting. IL-38 is notoriously unstable in biological assays^28^, and therefore identifying a robust assay was envisioned. Among the assays tested, IL-38 consistently suppressed cytokine production in two-way mixed lymphocyte reaction (MLR) assays using allogeneic lymphocyte populations from two different donors, which allows discovering immunotherapeutic effects on bi-directional T-cell activation^29^. Briefly, PBMCs from two donors were isolated and co-cultured for 6 days alone, or in the presence of IL-38, IL-1RA or IL-36RA. The latter were included to obtain information about a potential IL-38 receptor or differential mode of action. On day 6 after starting the MLR, we observed reduced levels of granzyme B (GrzB), IFN-γ, and TNF-α, but not IL-6 or IL-10 (Fig. 1A-B) in the supernatants of IL-38-treated versus control cells. IL-36RA only mildly decreased the secretion of TNF-α compared to control (Fig. 1A). Likewise, IL-1RA moderately suppressed the secretion of TNF-α and additionally reduced the concentration of IL- 10 in the supernatants (Fig. 1A). Thus, IL-38 potently suppressed inflammatory cytokine secretion during MLR, while acting independently of IL-1 and IL-36 receptors.

**Figure 1.**
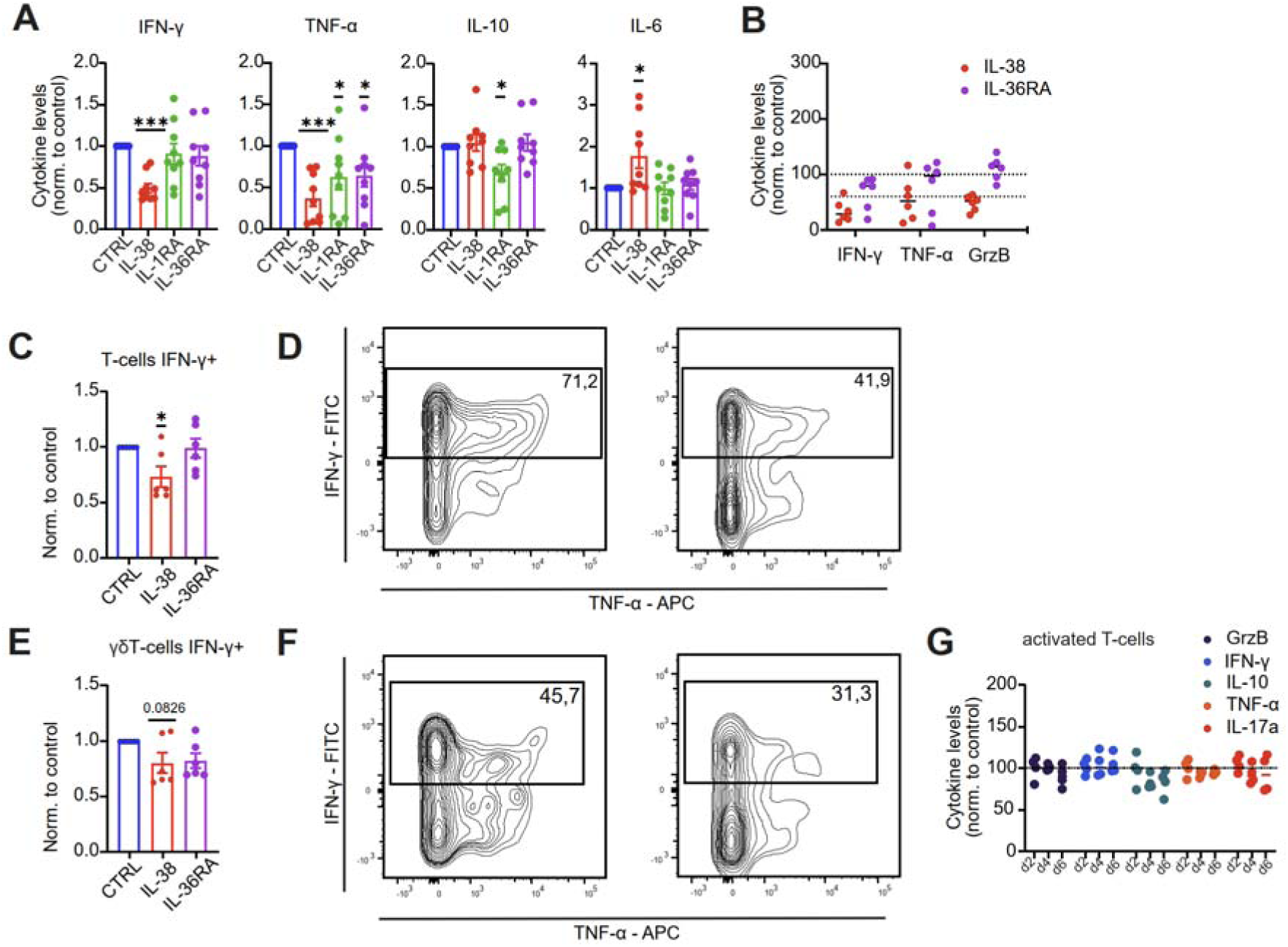
IL-38 inhibits inflammatory response in mixed lymphocyte reactions. Human PBMCs of unrelated donors were co-cultured for 6 days and treated with IL-38, IL-1RA, or IL-36RA (each 100 ng/ml). Levels of IFN-γ, TNF-α, IL-10, IL-6 **(A)** and Granzyme B (GrzB) **(B)** were measured on day 6 by Cytometric Bead Array. Cells were collected on day 6 and analyzed by intracellular FACS. Intracellular IFN-γ levels in total T-cells (**C–D**) and γδ T-cells (**E–F**) are shown. Normalized data from one representative experiment out of three. Each data point corresponds to one independent donor pair (n=6). (**G**) T-cells were cultured for 6 days in the presence of IL-38, and supernatants were harvested on days 2, 4, and 6 to evaluate cytokine concentrations. Normalized data of the IL-38-treated group compared to controls are shown. Data are from two independent experiments. Each data point corresponds to single donor (n=4). Data are shown as mean ± SEM. \**p* < 0.05, \*\**p* < 0.01, \*\*\**p* < 0.001, \*\*\*\**p* < 0.0001; *p*-values were calculated using one-sample *t*-test.

Importantly, endogenous IL-38 was not produced during MLR as determined by ELISA. To identify which cell populations were responsible for the reduction in cytokine secretion in response to IL-38, we performed intracellular flow cytometry on immune cell populations on day 6. Among lymphocytes, IFN-γ production was altered in αβ T-cells (Fig. 1C,D) and in γδ T-cells (Fig. 1E,F). Moreover, IL-38 had a minor impact on IFN-γ production by B cells, while IL-36RA increased IL-6 secretion (Suppl. Fig. 1A). To test whether IL-38 has a direct effect on T-cells, we isolated and activated T-cells with anti-CD3/CD28 beads alone or in the presence of IL-38. We did not observe any differences in cytokine secretion over a course of 6 days (Fig. 1G). Overall, these data indicate that IL-38 indirectly suppresses inflammatory and cytotoxic activities in T-cells in an allogeneic system and employs unique receptors or signaling pathways compared to other IL-1 family antagonists.

### IL-38 alters cell abundance and gene expression in mixed lymphocyte reactions

To identify which cell population conveys T cell suppression in the presence of IL-38, we performed longitudinal CITE-Seq of cells in MLR assays alone or in the presence of IL-38. Beforehand, we measured IFN-γ and TNF-α expression at the gene and protein levels daily over 6 days during the MLR to determine the starting point of the immune reaction. We observed increased cytokine secretion starting on day 3 and a first indication of the suppressive effect of IL-38 at this time point (Suppl. Fig.2A-C). We therefore collected cells on day 0, day 3 and day 5 during the MLR. 4 different donor combinations were pooled for CITE-Seq. After quality control and filtering of each replicate, we obtained 67455 cells. Principal component analysis of replicates per day and condition showed similar transcriptional profiles, demonstrating low batch-to-batch variation and supporting their pooling into a single dataset (Suppl Fig. 3A). Unsupervised Seurat clustering revealed 21 immune cell clusters (Fig. 2A). Based on expressions of cell-specific marker genes and proteins on the cell surface, we annotated 4 myeloid, 6 T-cell, 3 NKT-like cell, 3 B-cell, 2 NK-cell, basophil and miscellaneous clusters (Fig. 2A). Looking at immune cell abundance changes over time (Fig. 2A, Suppl. Fig. 3B-C ), we observed a similar reduction in the proportion of myeloid cells, basophils, NK-cells, NKT-like cells, γδ T-cells, naïve CD8+ T-cells and B cells, while CD4+ TEM, CD56^hi^ NK-cells and naïve activated B2-cells were expanded in both groups, which reflects the expected immunological responses. Surprisingly, we identified an increase in the percentage of both naïve CD4+ T-cell clusters, CD4+ TCM, NKT-like CD16^hi^ and NKT-like CD16^lo^ cells, CD8+ TEM, γδ T-cells and pDCs in the IL-38 group compared to control (Suppl Fig. 3C). In contrast, both naïve activated B-cell clusters and memory B-cells were decreased in the IL-38 group on day 3 and day 5 compared to control (Suppl Fig. 3C).

**Figure 2.**
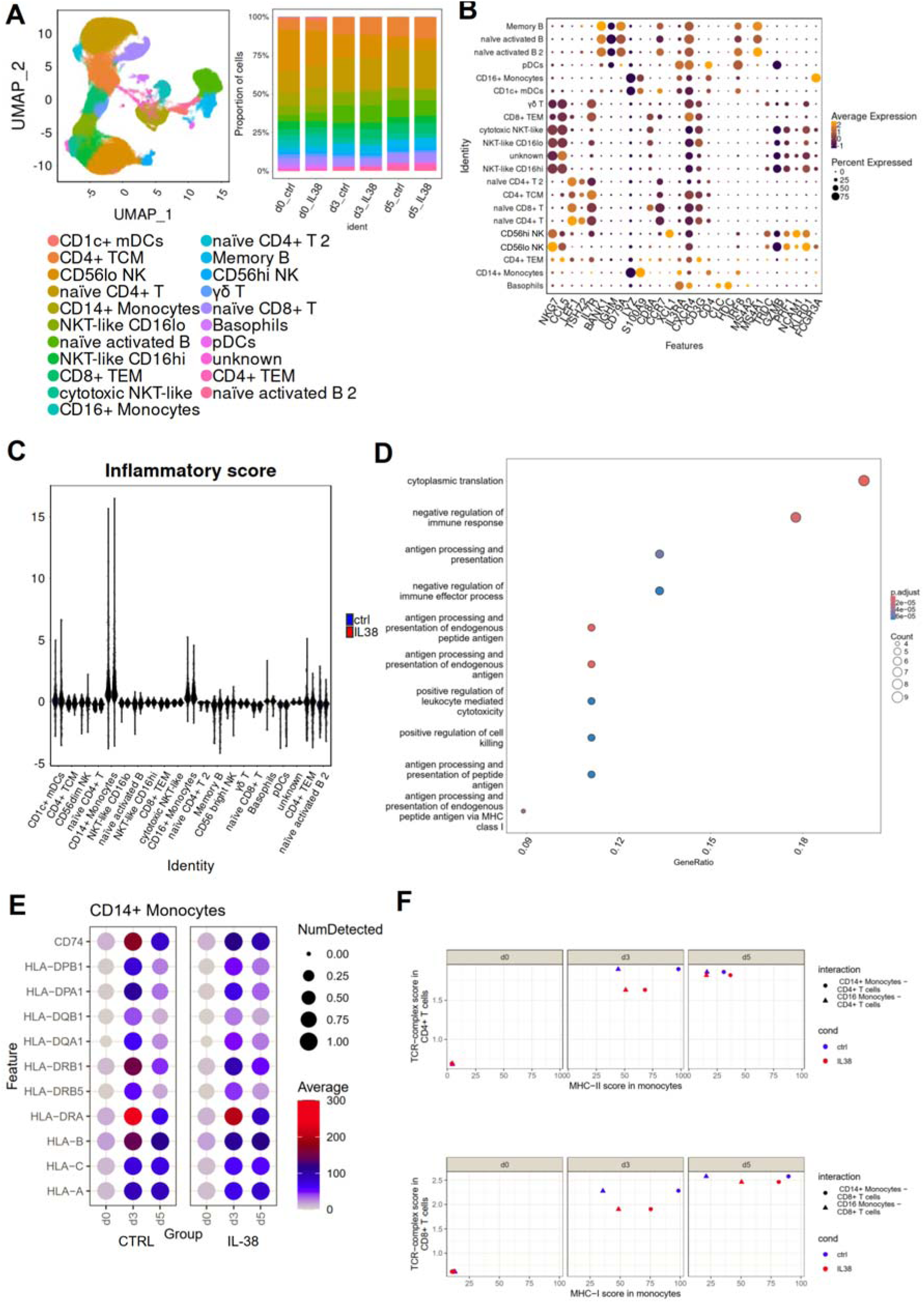
IL-38 alters immune cell populations during MLR. Human PBMCs from 4 unrelated donors were co-cultured in pairs and treated with IL-38 (100 ng/ml) daily. Cells were collected on days 0, 3, and 5 and analyzed by CITE seq. **(A)** A total of 21 cell clusters were detected in a unified UMAP plot (left panel), with their abundances shown by day and treatment condition (right panel). **(B)** Dot plot showing the annotation based on the main mRNA markers expressed in immune cell clusters. **(C)** Violin plots show the calculated Inflammatory score. **(D)** Gene Ontology (GO) analysis based on differentially expressed genes in CD14+ monocytes obtained on day 3. **(E)** Expression patterns of HLA-I and HLA-II related genes in CD14+ monocytes between control (left panel) and IL-38 (right panel) groups. **(F)** Receptor-ligand interaction model of CD4+ T-cells and CD14+/CD16+ monocytes (upper panel), and CD8+ T-cells and CD14+/CD16+ monocytes (lower panel) on day 3.

At the transcriptional level, various immune cell populations demonstrated altered gene expression in the presence of IL-38 compared to control (Suppl. Table 1). Major transcriptome changes were observed in CD14+ monocytes, both naïve CD4+ T-cell subsets, CD8+ T EM, both naïve activated B-cell subsets, NKT-like CD16^hi^ cells, γδ T-cells, NKT-like CD16^lo^ cells, CD56^lo^ NK-cells, memory B-cells, cytotoxic NKT-like cells, CD8+ naïve T-cells, and CD4+ CM T-cells. In addition, we calculated inflammation scores across all time points for all populations in our data set. Here, only myeloid cells (monocyte subsets and CD1c+ mDCs) showed an elevated score compared to other clusters (Fig. 2C). Of these myeloid populations, only CD14+ monocytes showed significant changes in gene expression on day 3 (Suppl. Table 1). Taking into consideration that the initial immune response in the MLR takes place on day 3 (Suppl. Fig. 2A-C), CD14+ monocytes appeared as the initial cellular target of IL-38.

### IL-38 decreases antigen-presentation capacity in monocytes

Focusing on CD14+ monocytes on day 3, differential gene expression and pathway analysis demonstrated that IL-38 diminished a significant number of genes involved in antigen processing and presentation, and regulation of the immune response and cytotoxicity pathways (Fig. 2D). Indeed, IL-38-treated CD14+ monocytes showed down-regulation of MHC-II, MHC-I and CD74 at the transcriptional level (Fig. 2E), which are key players in antigen presentation^2^. JAK2 and the primary myeloid transcription factor SPI1 were downregulated in IL-38-treated cells, suggesting a potential role for IL-38 in modulating myeloid cell fate (Suppl. Fig. 4A). Analysis of cell interaction revealed elevated interaction between CD4+ T-cells and monocyte populations on day 3 (Fig. 2F), which was diminished in the IL-38-treated group. On day 5 there were no differences in interaction scores. Similar results were observed when calculating interactions between CD8+ T-cells and CD14+ and CD16+ monocytes based on TCR-complex and MHC-I scores (Fig. 2 F). The effect was likely due to an impact on monocytes, since expression of TCR-complexes was largely unaltered (Suppl. Fig. 4B-C). As interaction with CD4+ T cells is specific for antigen-presenting cells, we further focused on CD4+ T cell populations. We calculated Th1/APC interaction, activation and checkpoint expression scores in all CD4 T-cell subsets present in our data set (Suppl. Fig. 4D-E). IL-38 diminished the Th1 score in all CD4+ T-cell populations on day 3 and day 5. Surprisingly, checkpoint expression and activation scores were dramatically altered only in CD4+ EM T-cells on day 3 and day 5 (Suppl. Fig. 4F-G).

Subsequent pathway analysis in CD4+ T-cells demonstrated that T-cell differentiation and T-cell selection pathways were altered in CD4+ CM T-cells (Suppl. Fig. 5A), and T-cell differentiation and T-cell activation were altered in CD4+ naive T-cells on day 3 (Suppl. Fig. 5B). Thus, IL-38 downregulates antigen-presentation related genes in CD14+ monocytes in the initial phase of an allogeneic reaction, which may affect differentiation and activation of CD4+ T-cells.

### IL-38 alters MHC-II expression during monocyte to macrophage differentiation

IL-38 affected genes related to antigen presentation in monocytes on day 3 in the MLR. Therefore, we tested whether IL-38 affects MHC-II expression on isolated human classical monocytes alone (control) or in the presence of LPS or TNF-α *in n vitro*. These stimuli were chosen since they are the major mediators of GvHD in the first phase^6^. No significant changes in HLA-DR and CD86 surface expression were observed (Fig. 3A). In the MLR, monocytes differentiate into macrophages over time. Therefore, we next determined whether IL-38 has an impact on human macrophage differentiation *in vitro*. Human primary monocytes were differentiated to macrophages in the presence of human plasma for 7 days. Daily IL-38 treatment transiently decreased HLA-DR expression on day 3 as determined by flow cytometry, while its expression was restored on fully differentiated macrophages (Fig. 3B). CD86 expression was not significantly affected in this setting (Fig. 3B), and among other macrophage polarization markers, only CD206 was decreased in the presence of IL-38 on day 7 (Suppl. Fig. 6A). The impact of IL-38 on differentiating monocytes was more pronounced when we used purified CD14+ cells differentiated in the presence of M-CSF. Expression of HLA-DR and CD86 on day 3 were decreased but largely re-established on day 7 (Fig. 3C-D). Thus, IL-38 diminishes HLA-DR surface expression transiently during monocyte-to-macrophage differentiation. This observation is substantiated by diminished expression of the myeloid transcriptional regulators SPI1 and CIITA in macrophages on day 3 upon IL-38 treatment (Fig. 3E), which was also observed in the CITE-seq data (Fig. 3F). When taking a closer look at gene expression levels between day 3 and day 5 in control and IL-38 groups in CD14+ monocytes in CITE-seq dataset, GO analysis revealed that cells in both groups expressed genes involved in myeloid cell differentiation (Suppl. Fig. 6B-C). However, only IL-38-treated cells demonstrated alterations in genes involved in leukocyte activation and migration, lymphocyte activation and phagocytosis (Suppl. Fig. 6B). Importantly, treatment of differentiating macrophages with IL-38 only once on day 3 did not affect HLA-DR and CD86 surface expression, compared to daily treatment with IL-38, indicating a cumulative effect (Fig. 3G). To determine if changes in MHC-II expression translate into altered antigen presentation capacity, we performed early macrophage/T-cell co-cultures in allogeneic and autologous settings, with or without the presence of model antigens and IL-38 (Fig. 3H-I). For that, monocytes were differentiated to macrophages for 3 days with or without addition of IL-38. Antigens were added on day 2 and allogeneic or autologous T cells were added, followed by co-culture for another 3 days (Suppl. Fig. 7). IL-38 completely prevented the proliferation of T-cells triggered by antigens in an allogeneic (Fig. 3H), but not in an autologous setting (Fig. 3I). Overall, these data show that IL-38 transiently alters the differentiation of macrophages to limit antigen presentation capacity and T-cell activation during monocyte-to-macrophage differentiation.

**Figure 3.**
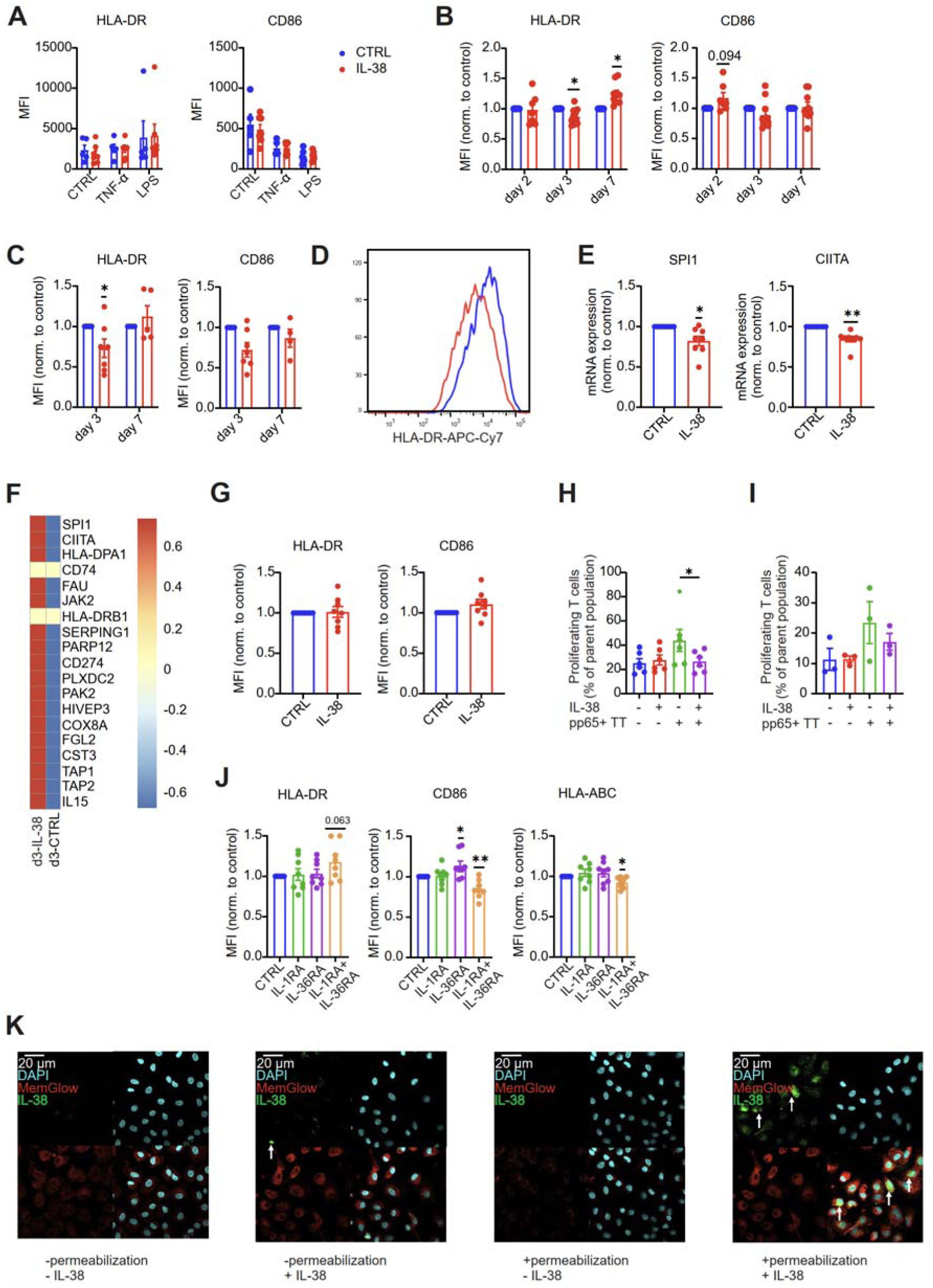
IL-38 transiently modulates MHC-II surface expression and antigen-presentation capacity during macrophage differentiation. Human CD14+ monocytes were cultured for 24 hours, unstimulated (CTRL) or stimulated with TNF-α (10 ng/ml), or with LPS (10 ng/ml), and treated with IL-38 (100 ng/ml). The surface expressions of HLA-DR (left panel) and CD86 (right panel) were determined by flow cytometry **(A)**. Each data point represents an individual donor (n=5-7), and the data are from 2 independent experiments. **(B)** Monocytes were differentiated in the presence of human plasma for up to 7 days and treated daily with IL-38 (100 ng/ml). The surface expressions of HLA-DR (left panel) and CD86 (right panel) on days 2, 3, and 7 were determined by flow cytometry. Data are means ± SEM. Each data point represents an individual donor (n=7-11), and the data are from 3 independent experiments. **(C)** Monocytes were differentiated in the presence of rhM-CSF (50 ng/ml) for up to 7 days and treated daily with IL-38 (100 ng/ml). The surface expressions of HLA-DR (left panel) and CD86 (right panel) on days 3 and 7 were determined by flow cytometry. Each data point represents an individual donor (n=5-7), and the data are from 2 independent experiments. **(D)** A representative histogram of HLA-DR surface expression on day 3 is shown. **(E)** Early macrophages were differentiated in the presence of rhM-CSF (50 ng/ml) and collected on day 3. SPI1 (left panel) and CIITA (right panel) mRNA expression were determined by qPCR. Each data point represents an individual donor (n=7), and the data are from 3 independent experiments. **(F)** Representative heatmap of average expression of down-regulated genes in CD14+ monocytes on day 3 (control and IL-38) from CITE-seq data is shown. **(G)** Early macrophages were cultured in the presence of human plasma and treated with IL-38 (100 ng/ml) once on day 3. Surface expression of HLA-DR (left) and CD86 (right) determined by flow cytometry. Each data point represents an individual donor (n=8), and the data are from 2 independent experiments. **(H, I)** Macrophages were treated with IL-38 daily (100 ng/ml), on day 2, cells were pulsed with Tetanus Toxoid (TT) at 1 µg/ml and pp65CMV peptivator (pp65) at 1 µg/ml of each peptide, and co-cultured with allogeneic **(H)** or autologous **(I)** T-cells for up to 3 days at a M:T-cell ratio of 1:2. T-cell proliferation was determined by cell proliferation dye eFluor670 dilution via flow cytometry. Each data point represents an individual donor (**H**: n=6, **I**: n=3). **(J)** Macrophages were cultured in the presence of human plasma for up to 3 days and treated daily with IL-1RA (100 ng/ml), IL-36RA (100 ng/ml), or the combination of both. Surface expressions of HLA-DR, CD86, and HLA-ABC Cells were determined by flow cytometry. Each data point represents an individual donor (n=8), and the data are from 2 independent experiments. **(K)** Early macrophages were cultured for up to 3 days in the presence of rhM-CSF (50 ng/ml) and treated with IL-38 (10 µg/ml) for 1h. Immunocytochemistry images show early macrophages stained with anti-IL-38 (green), MemGlow (red), and DAPI (cyan). Scale bars represent 20 μm. Representative images of one individual donor out of 3 independent experiments are shown. All data are means ± SEM and are biological replicates. ^∗^p < 0.05; ^∗∗^p < 0.01; ^∗∗∗^p < 0.001; p values were calculated using a two-way ANOVA with Tukey’s multiple-comparisons correction (**A**), one-sample *t-* test for normalized data (**B, C, E, G, J**), and one-way ANOVA with Dunn’s multiple-comparisons correction (**H**).

### IL-38 affects early macrophages through a receptor-independent pathway

Next, we examined mechanisms underlying the IL-38-mediated effect on early macrophage differentiation, starting with testing the involvement of IL-1 family receptors. Macrophages showed moderate expression of IL-1R1, while IL-1RAPL1 and IL-36R were not or barely expressed (Suppl. Fig. 8A). We analyzed if daily treatment with IL-1RA, IL-36RA, or with a mix of both cytokines had a similar effect on early macrophage differentiation as IL-38. HLA-DR surface levels were unchanged in cells treated with IL-1RA, IL-36RA or the combination of both (Fig. 3J), while CD86 was elevated in IL-36RA-treated cells, but diminished in the combination group compared to control (Fig. 3J). HLA-ABC surface level decreased in the presence of both IL-1RA and IL-36RA (Fig. 3J). These data suggest that IL-38 likely employs a receptor-independent mechanism to regulate HLA-DR and CD86 expression. IL-38 has been postulated to have intracellular functions as well ^28^. Therefore, we examined a potential uptake of exogenous IL-38 into macrophages. We treated macrophages with 10 µg/ml of IL-38 followed by fluorescence confocal imaging. Untreated macrophages or IL-38-treated but unpermeabilized cells did not show IL-38 expression (Fig. 3K). However, IL-38-treated and permeabilized cells showed a bright IL-38 signal, indicating that exogenous IL-38 was taken up by macrophages (Fig. 3K). Mechanistically, intracellular IL-38 was postulated to affect metabolism and/or ROS expression^30^. However, in human macrophages, we observed neither an altered oxygen consumption rate (Suppl. Fig. 8B) nor an impact on ROS production (Suppl. Fig. 8C) determined by Seahorse extracellular flux analysis and flow cytometry, respectively. These data indicate that IL-38 transiently affects MHC-II cell surface expression during macrophage differentiation by a so-far unappreciated, receptor-independent mechanism.

### IL-38 ameliorates murine GvHD

Given the immunomodulatory effects of IL-38 in the allogeneic human *in vitro* system, we sought to confirm these findings in a murine model of GvHD. We used a mouse model of complete MHC-mismatched bone marrow transplantation (allo-BMT) using T cell-depleted bone marrow followed by an allogeneic T-cell transfer. These donor T-cells contained a Nur77-GFP reporter system to track T cell receptor (TCR)-mediated T-cell activation *in vivo*^31^. Mice received IL-38 3 times a week at a concentration of 1 mg/kg (Fig. 4A) starting on day 3. IL-38 treatment significantly improved survival (Fig. 4B) and disease scores in mice (Fig. 4C). However, there were no changes in weight loss between control and IL-38 groups (Suppl. Fig 9A). These data suggest a potential prophylactic effect of IL-38 in murine GvHD. When performing flow cytometric immune profiling of murine blood (Suppl. Fig. 10), we did not identify differences in major blood donor-derived immune cell populations between control and IL-38 groups (Suppl. Fig. 9B). The percentage of engrafted (H2-kb) cells and proportions of donor-derived CD4+ T-cells and CD8+ T cells were similar (Suppl. Fig 9C-D) in control and IL-38 groups. Graft-derived CD8+ eff/CD8+ mem and CD8+ naïve T-cells remained unchanged between control and IL-38 groups (Fig. 4D). However, we observed a reduction in the numbers of graft-derived memory CD4+ T-cells and an expansion of Tregs (Fig. 4E-F) in the IL-38 group compared to control. In addition, we found a reduction of donor-derived GFP+ CD4+, but not of GFP+ CD8+ T-cells, indicating a major difference in TCR engagement in helper T-cells between treatment groups in the blood (Fig. 4G-H), which could be explained by diminished TCR-MHC-II interaction in the IL-38-treated group, as observed in the human MLR assays. Cytokine concentrations (IFN-γ, TNF-α, IL-6 IL-17a, IL-10, MCP-1 and IL-1β) were similar in murine serum between groups at the endpoint (Suppl. Fig. 9E). Immune cell profiles of engrafted (H2-kb) (Suppl. Fig. 11A), donor-derived immune cells (Suppl. Fig. 11B), and proportions of donor-derived CD4+ T-cells and CD8+ T-cells were unchanged (Suppl. Fig. 11C) in the spleen. Analysis of subpopulations of graft CD4+ T-cells and graft CD8+ T-cells revealed no significant differences between IL-38-treated and control animals (Fig. 4I-J).

**Figure 4.**
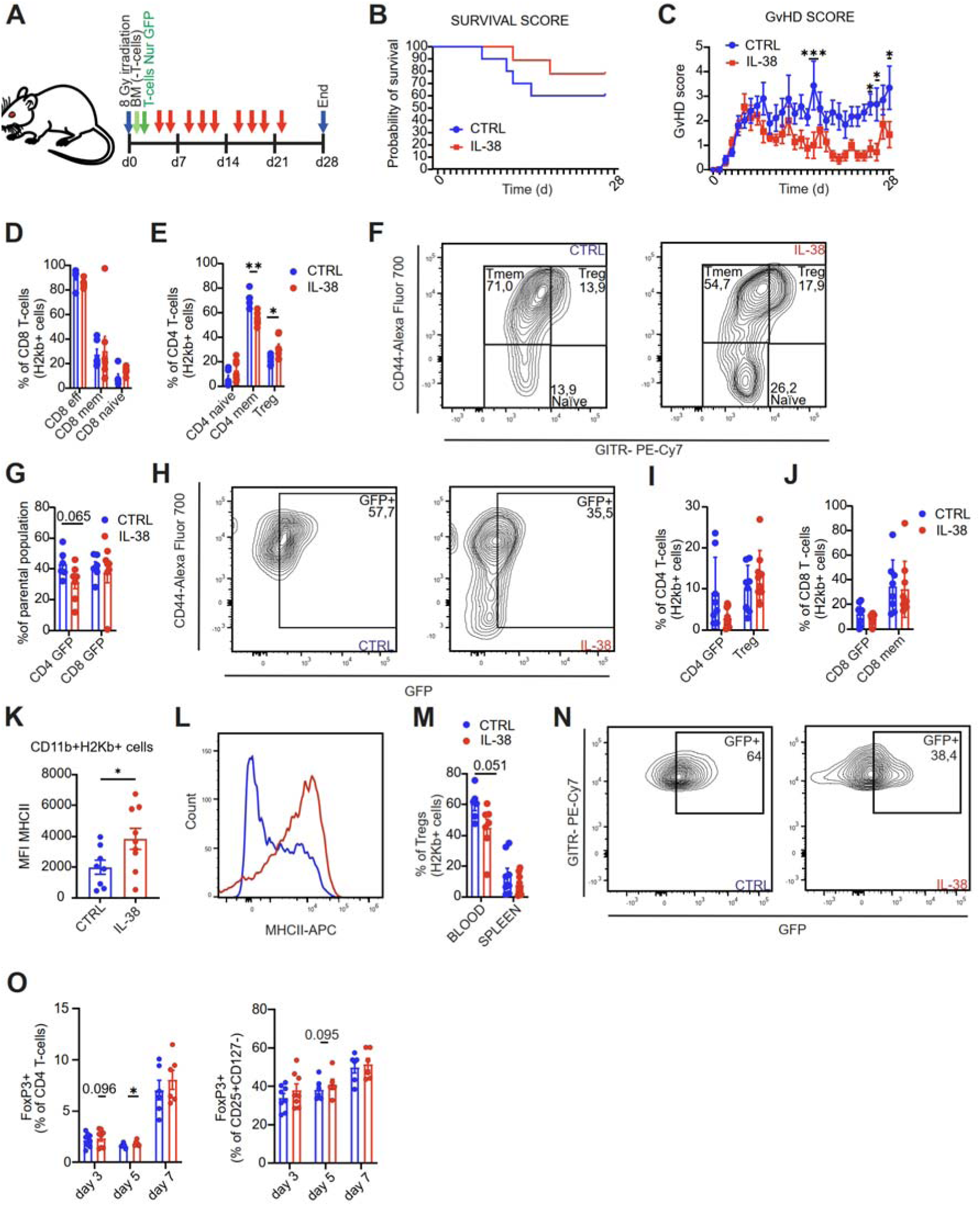
IL-38 ameliorates murine GvHD. **(A)** The cartoon displays the experimental setup. BALB/c mice (host) were irradiated on day 0 and received T-cell-depleted bone marrow (BM) cells from wild type (wt) C57BL/6 and T-cells from Nur77^GFP^ mice on the following days. IL-38 was given (i.p) three times a week. **(B)** Survival curves and **(C)** GvHD disease scores of mice treated with IL-38 or vehicle (n=10 control, n=9 IL-38). **(D)** The graph indicates the percentage of CD8 T-cell subsets of donor-derived cells in the blood of CTRL and IL-38-treated mice. Data are means ± SEM (n=6). **(E,F)** The graph **(E)** and representative FACS plot **(F)** show percentage of CD4 T-cell subsets of donor-derived cells in blood of CTRL and IL-38 treated mice. Data are means ± SEM (n=6). **(G,H)** The graph **(G)** and representative FACS plot **(H)** show the percentage of GFP+ cells of the parental population (CD4+ T-cells) of donor-derived cells in the blood of CTRL and IL-38 treated mice. Data are means ± SEM (n=6). **(I,J)** Relative proportion of CD4+ T-cell **(I)** and CD8 T-cell **(J)** subsets of donor-derived cells in the spleen of CTRL and IL-38 treated mice. Data are means ± SEM (n=8 control, n=9 IL-38). **(K,L)** The graph **(K)** and representative histogram **(L)** show MHC-II surface expression of donor-derived APCs (CD11b+H2kb+) in spleens of control and IL-38-treated mice. Data are means ± SEM (n=8 control, n=9 IL-38). **(M,N)** The graph **(M)** and representative FACS plot **(N)** show the percentage of GFP+ cells within the Treg population of donor-derived cells in blood and spleen. Data are means ± SEM (n=6-7). **(O)** Two-way MLR with human PBMCs was performed. PBMSc form anonymous donors were co-cultured in pairs for up to 7 days. The IL-38 treatment was added daily at a concentration of 100 ng/ml. Cells were collected on days 3, 5 and 7 and percentages of FoxP3+ cells of CD4+ T-cells (left panel) and of CD25+ CD127 CD4+ T-cells are shown. Normalized data compared to control. Data are means ± SEM (n=6) of 3 independent experiments. All data are biological replicates. ^∗^p < 0.05; ^∗∗^p < 0.01; ^∗∗∗^p < 0.001; p values were calculated using multiple unpaired *t-* test (**D-E, I-J**), unpaired *t*-test (**G, K, M**) or one-sample *t*-test (**O**).

Notably, donor-derived APCs (CD11b+H2Kb+ cells) showed increased MHC-II surface expression in the IL-38 group compared to control (Fig. 4K-L). We did not observe significant changes in MHC-II surface expression in host-derived APCs (CD11b+H2Kd+) (Suppl. Fig. 11D). Given the expansion of Tregs in the blood of IL-38-treated animals, we examined whether these cells were generated upon TCR engagement. Surprisingly, the number of GFP+ Tregs in blood, but not in spleen, decreased (Fig. 4M-N), suggesting a TCR-independent mechanism. To reconfirm an effect in Treg generation upon IL-38 treatment, we revisited the MLR data. Here, IL-38 indeed increased FoxP3+ T-cells on day 5 (Fig. 4O). Collectively, these data provide evidence that IL-38 exerts immunosuppressive effects on graft immunity at the systemic level, which includes the expansion of Tregs, likely in a TCR-independent manner.

### IL-38 diminishes inflammation in the periphery in GvHD

After evaluating systemic inflammation, we determined whether IL-38 has an impact on the key target organs in GvHD, namely, the skin, liver, and colon^32–34^. Animals in both groups did not show visible signs of skin inflammation. These observations were also supported by immunohistochemical examination of skin samples (data not shown). However, since IL-38 is dominantly expressed in the skin and proposed to maintain homeostasis in keratinocytes^28^, we analyzed publicly available single-cell transcriptome data from^35^ towards IL-38 mRNA levels in skin of healthy individuals compared to skin samples of patients with GvHD. IL-38 (IL1F10) was only expressed in differentiated keratinocytes of healthy individuals, but its expression was lost in GvHD (Suppl. Fig. 12A-B). Thus, a lack of IL-38 correlates with cutaneous GvHD. Flow cytometric immune profiling of liver cells (Suppl. Fig. 13) showed no difference in grafted cells in liver samples between control and IL-38 groups (Suppl. Fig. 14A). There were no differences in major immune cell populations (Suppl. Fig. 14B). The proportions of donor-derived CD4+ and CD8+ T-cells (Suppl. Fig. 14C) were unchanged between groups. In addition, we did not identify significant differences in MHC-II expression levels on APCs (Suppl. Fig. 14D). Graft-derived CD8+ T-cell subsets remained unchanged between groups (Fig. 5A). However, graft Tregs were again expanded in the IL-38 group compared to the control (Fig. 5B-C). There were no significant differences in GFP+ CD4+ and CD8+ T-cells. In contrast, the immuno-histochemical examination of the liver samples showed a significant decrease in activated macrophages (MHC-II+ F4/80+) and CD4+ T-cells (Fig. 5D-E), albeit no differences in Treg numbers in the IL-38 group compared to control. For colon, we examined both intraepithelial lymphocytes (IEL) and lamina propria lymphocytes (LPL) fractions. The percentage of donor-derived immune cells in LPL was similar in control and IL-38 groups (Suppl. Fig. 15A). Flow cytometric immune profiling of graft-derived (CD45+ H2Kb+ cells) and host-derived cells was performed (Suppl. Fig. 16), revealing no differences between control and IL-38-treated animals (Suppl. Fig. 15B-C). The proportions of donor-derived CD4+ T- and CD8+ T- cells remained unchanged between groups (Suppl. Fig. 15D). There were no differences in GFP+ CD4+ and CD8+ T-cells, CD4+ T-cells, and CD8+ memory T-cells between control and IL-38 groups in the LPL (Fig. 5F-G). However, graft-derived Tregs were expanded similarly to the observations in the blood (Fig. 5G-H), together with augmented MHC-II surface expression on donor-derived APCs (CD11b+ H2Kb+ cells) (Fig. 5I-J). Accordingly, IHC evaluation of the LPL compartment suggested elevated numbers of MHC-II+ macrophages (MHC-II+ F4/80+), CD4+ Tregs (CD3+ CD4+ FoxP3+), and CD8+ Tregs (CD3+ CD8+ FoxP3+), as well as a decreased proportion of total macrophages (F4/80+) (Fig. 5K-L). Immune cell distribution in IEL was comparable between groups (data not shown). These cells were fully donor-derived (Suppl. Fig. 15E). Immuno-profiling of all intraepithelial immune cells (Suppl. Fig. 17) also revealed increased Tregs in the IL-38-treated group compared to control (Fig. 5M-O; Suppl. Fig. 15F). Other donor-derived T-cell subpopulations were unchanged (Fig. 5M-N; Suppl. Fig. 15F-G) except for decreased CD4+ mem T-cells together with expanded Tregs (Fig. 5M-O). GFP+ CD8+ and CD4+ T-cells were comparable between groups (Fig. 5M-N). There was no difference in MHC-II surface expression in host-derived epithelial cells (Suppl. Fig. 15H). These observations were partly supported by IHC measurements, where the numbers of CD4+ Tregs (CD3+CD4+FoxP3+), CD8+ Tregs (CD3+CD8+FoxP3+), and MHC-II+ epithelial cells were increased, and macrophages (F4/80+) were elevated in the IEL of the IL-38-treated group (Fig. 5P-Q). These data indicate that IL-38 limits gut inflammation during GvHD, as evidenced by an expansion of donor-derived Tregs.

**Figure 5.**
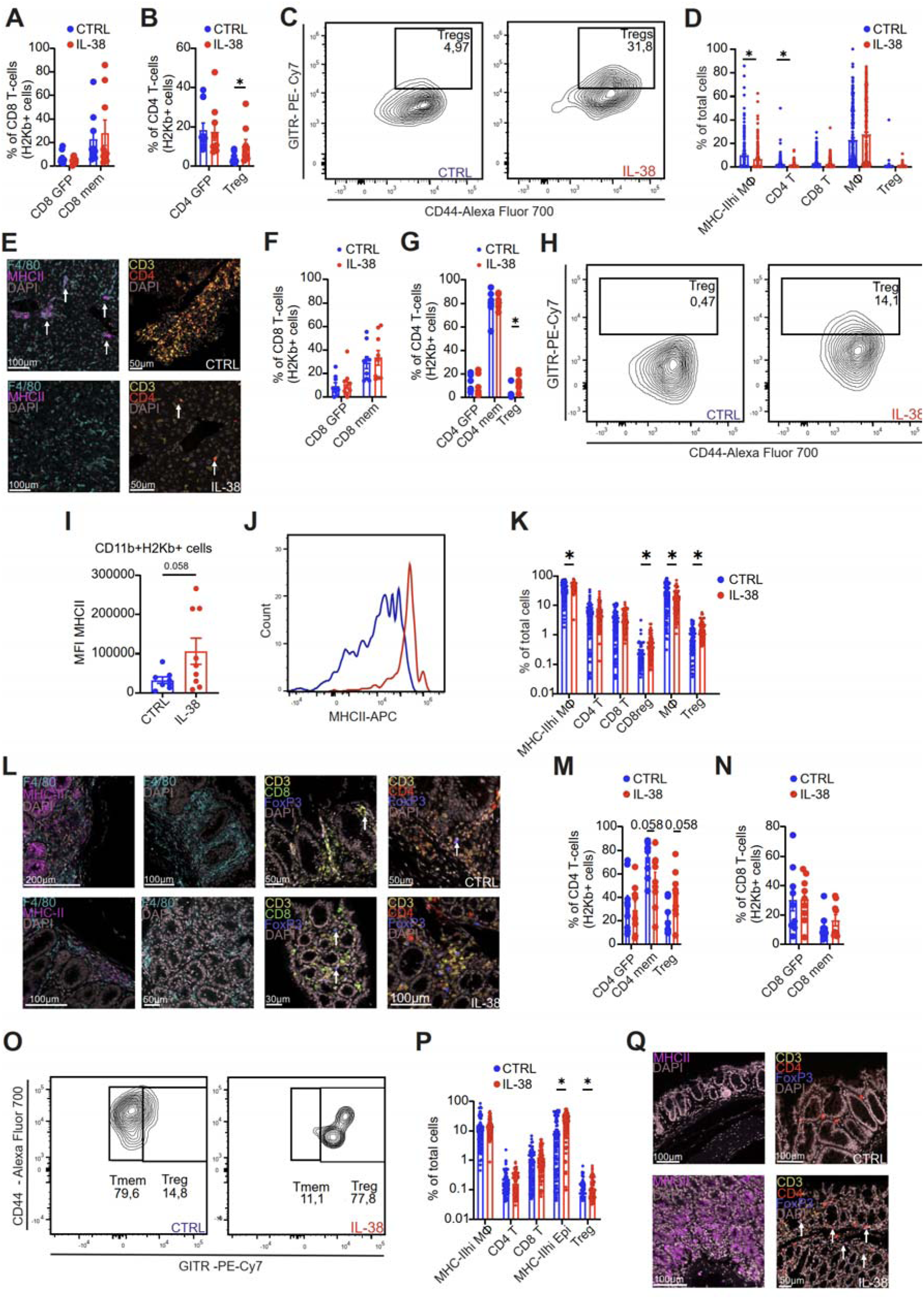
IL-38 alleviates inflammation in liver and intestines during GvHD. BALB/c mice (host) were irradiated on day 0 and received T-cell-depleted bone marrow (BM) cells from wt C57BL/6 and T-cells from Nur77^GFP^ mice on the following days. IL-38 was given (i.p) three times a week. **(A)** CD8+ T-cell proportions of donor-derived cells in the liver determined by flow cytometry. Data are means ± SEM (n=9). **(B,C)** The graph **(B)** and representative FACS plot **(C)** of CD4+ T- cell subsets of donor-derived cells in the liver determined by flow cytometry. Data are means ± SEM (n=9). **(D,E)** The graph **(D)** and representative images **(E)** of immune cells in the liver determined by immunohistochemistry stained with anti-F4/80 (cyan), anti-MHC-II (magenta), anti-CD3 (yellow), anti-CD4(red), and DAPI (grey). The control group is on the upper panel, and IL-38 in the bottom panel. Data are means ± SEM (n=8-9). Scale bars represent 100 μm or 50 μm. **(F)** CD8+ T-cell proportions of donor-derived cells in the liver determined by flow cytometry. Data are means ± SEM (n=8-9). **(G, H)** Graph of CD4+ T-cell subsets **(H)** and representative FACS plot **(H)** of Tregs of donor-derived cells in the LPL determined by flow cytometry. Data are means ± SEM (n=8-9). **(I, J)** The graph **(I)** and representative histogram **(J)** show MHC-II surface expression of donor-derived APCs (CD11b+H2kb+) from LPL in control and IL-38-treated mice determined by flow cytometry. Data are means ± SEM (n=8-9). **(K, L)** The graph **(K)** and representative images **(L)** of immune cells in the LPL determined by immunohistochemistry stained with anti-F4/80 (cyan), anti-MHC-II (magenta), anti-CD3 (yellow), anti-CD8 (green), anti-CD4(red), anti-FoxP3 (blue), and DAPI (grey). Scale bars represent 100 µm and 200 µm. Data are means ± SEM (n=8-9). **(M, N)** The graph **(M)** of CD4+ T-cell and **(N)** of CD8+ T-cell proportions of donor-derived cells in the IEL compartment determined by flow cytometry. Data are means ± SEM (n=9). **(O)** Representative FACS plot of Tmem and Tregs subpopulations of donor-derived CD4+ T-cells in the IEL compartment determined by flow cytometry. **(P,Q)** Graph **(P)** and representative images **(Q)** of immune cells in the IEL determined by immunohistochemistry stained with anti-MHC-II (magenta), anti-CD3 (yellow), anti-CD4(red), anti-FoxP3 (blue), and DAPI (grey). Scale bars represent 100 µm or 200 µm. Data are means ± SEM (n=8-9). All data are biological replicates (individual animals). ^∗^p < 0.05; ^∗∗^p < 0.01; ^∗∗∗^p < 0.001; p values were calculated using multiple unpaired *t-* test **(A, B, D, F, G, K, M, N, P)**, or unpaired *t*-test **(I)**.

### IL-38 limits inflammation in a human intestinal GvHD model

Data from the murine GvHD model indicated a protective role of IL-38 particularly in gut inflammation. To obtain insights into a human setting, we analyzed GvHD-like inflammation in a colon organ-on-chip system that allows determining immune cell infiltration across an endothelial barrier under flow conditions and damage to the epithelial compartment. In this setting, IL-38 attenuated the concentration of pro-inflammatory mediators such as IL-6 and Granzyme B (Fig. 6A). We did not detect significant differences in the PBMC infiltration ratio and epithelial volume and barrier disruption between the control and IL-38 groups (Fig. 6B-D), even though epithelial volume and barrier were significantly disturbed when PBMCs were infiltrated into the organ-chips compared to organ-chips without PBMCs, while changes were not significant when IL-38 was added together with PBMCs. The proportion of T cells in the epithelial layer decreased in the IL-38 group compared to the control. There were no significant differences in migrated individual T-cell subsets and myeloid cells to the epithelial layer between groups (Fig. 6E). Thus, IL-38 also limited aspects of inflammation in a human intestinal GvHD model.

**Figure 6.**
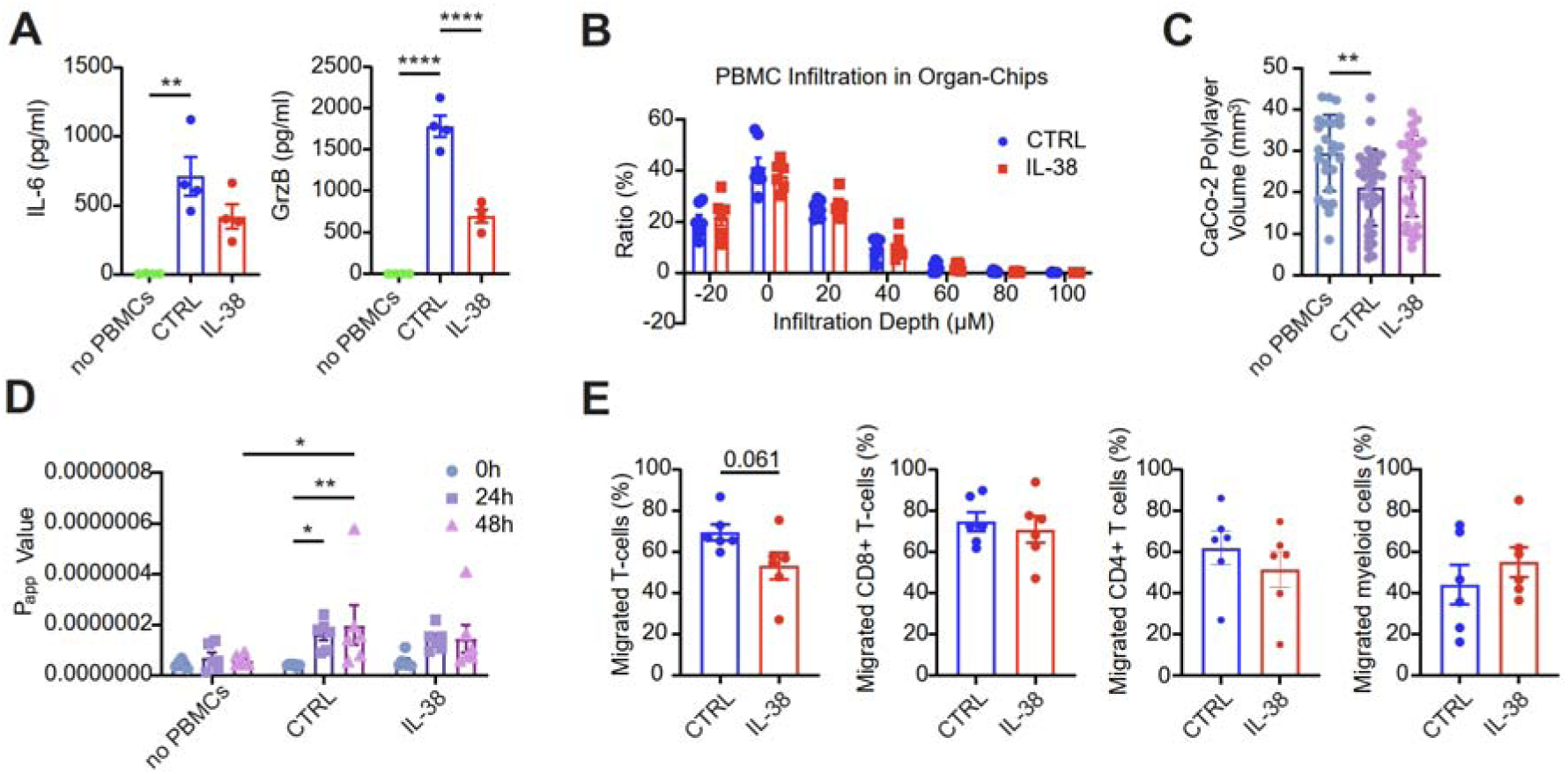
IL-38 inhibits inflammatory response in a human intestinal organ-on-chip model. Caco-2 cells and HUVECs were seeded into the apical epithelial channel and the basal endothelial channel of Emulate Colon Intestine-Chips, respectively. The cells were cultured in organ-chips for 7 days. PBMCs were pre-primed in the presence of Caco-2 cells for 4 days. On day 8, the organ-chips were treated with TNF-α to induce membrane disruption. PBMCs were then introduced and maintained in the organ-chips for up to 72 h. During this 72-h period, IL-38 (100 ng/mL) was administered daily. **(A)** Concentrations of IL-6 (left panel) and Granzyme B (right panel) were determined in supernatants from organ-chip systems at the endpoint (72 h post PBMC administration). Data are means ± SEM (n=4) of 3 independent experiments. **(B)** Ratio of infiltrated cells in both groups quantified by confocal microscopy is shown. Data are means ± SEM (n=6-7) of 3 independent experiments. **(C)** The volume of the epithelial layer was quantified by confocal microscopy. Data are means ± SEM (n=6-7) of 3 independent experiments, with individual regions of interest taken as data points. **(D)** Apparent Permeability Coefficient (P_app_) across the epithelial and endothelial barrier after PBMC administration with or without addition of IL-38. Data are means ± SEM (n=6-7) of 3 independent experiments. **(E)** Cells were collected at the endpoint from both channels, analyzed by flow cytometry, and migration rate was determined. Data are means ± SEM (n= 6) of 3 independent experiments. All data are biological replicates. ^∗^p < 0.05; ^∗∗^p < 0.01; ^∗∗∗^p < 0.001; p values were calculated using one-way ANOVA **(A,C)**, mixed-effects model (REML) **(B)**, two-way ANOVA followed by Dunnett’s multiple comparisons test **(D)** or unpaired *t*-test **(E)**.

### Materials and Methods GvHD mouse model

An allogeneic transplantation model with differences in major histocompatibility (MHC) loci between Balb/c recipient mice (MHC H2^d^) and C57BL/6 donor mice (MHC H2^b^) was utilized to trigger acute GvHD-like inflammatory reactions. Balb/c mice, Nur77-GFP C57BL/6 mice (JAX stock #040426) and wild type C57BL/6 mice were obtained from the Jackson Laboratory and housed under specific pathogen-free (SPF) conditions. Balb/c mice were lethally irradiated with 8 Gy using a cesium radiation device (day 0) and each recipient mouse received two sequential intravenous tail vein injections of T-cell-depleted bone barrow cells (5 * 10e6 cells) from C57BL/6 mice after 24 h (day 1) and splenic enriched T cells (0.5 * 10e6 cells) from Nur77-GFP C57BL/6 mice after 48 h (day 2). The cells were injected in a total volume of 100 µl sterile PBS under short-term isoflurane anesthesia. The antibiotic Baytril (15 mg/kg) was applied to drinking water and renewed weekly throughout the experiment to prevent infection. From the 2^nd^ day after irradiation, mice received i.p. injection of either the vehicle (control), or IL-38 (1 mg/kg;3 times a week) for a time span of 21 days. GvHD disease severity score was determined daily based on weight loss, posture, activity, fur texture, skin integrity and stool consistency. At the end of the experiment, mice were euthanized after i.p. injection of ketamine (180 mg/kg) and xylazine (10 mg/kg) followed by cardiac perfusion with 0.9% saline solution. Blood, spleen, liver, and colon samples were thereafter collected for follow-up analyses. All animal experiments were approved by, registered at, and followed the guidelines of the Hessian animal care and use committee (FU/2103).

### Human organ-chip model of intestinal GvHD

CaCo-2 cells were cultured in Advanced DMEM (Gibco) with GlutaMAX, 20% FCS, 1% Penicillin-Streptomycin, 0.1% Gentamicin, and HUVEC cells in endothelial growth medium (Pelobiotech) with 2% FCS, 1% Penicillin-Streptomycin, 0.1% Gentamicin. Following the manufacturer’s protocol for a colon intestine-chip from Emulate, the CaCo-2 cells and HUVEC cells were seeded, respectively, into the apical epithelial channel and the basal endothelial channel. Organ-chips, sequentially connected to pods and the Zoe culture module (all from Emulate), were cultured under conditions including cyclic stretches and continuous media flow for a week, allowing for cell proliferation and polarization of CaCo-2 cells into 3D epithelium. PBMCs were immune-primed by co-culture with CaCo-2 monolayers for four days in RPMI 1640 with 3% human plasma, 1% Penicillin-Streptomycin, 2% essential amino acids, 1% non-essential amino acids, 1 mM sodium pyruvate, 10 mM HEPES, CD3/CD28 T cell activation antibodies, while 10 ng/ml IL-2 (PeproTech), 10 ng/ml IL-7 (PeproTech) and 5 ng/ml IL-15 (PeproTech) and 50 µM β-mercaptoethanol (Pan Biotech) were applied to the co-culture and supplemented every two days. Measurement of cytokine release (IL-1β, IL-6 and IL-17α) in the co-culture supernatant via cytometric bead array (CBA) was performed to evaluate priming-induced immunity. On the 9^th^ day, after a 24-hour pre-treatment with 50 ng/ml TNF-α within the endothelium channel inducing endothelial permeability, primed PBMCs were collected following a dead cell removal protocol and labeled with a membrane marker PKH26 (Sigma-Aldrich), enabling fluorescence microscopy afterwards. PBMCs were infiltrated into the endothelium channel for 4h. The established intestinal GvHD organ-chips were maintained for 3 days under stretch and flow conditions, with or without addition of 100 ng/ml IL-38.

At the endpoint (72 h post PBMC administration), cells in each channel were stained with Hoechst (BD Biosciences) followed by confocal microscopy to assess the epithelium 3D structure and PBMC infiltration depth. Images were acquired on the Stellaris confocal platform with a 10X objective. The total volume of Hoechst in the epithelium channel was calculated on Fiji ImageJ, as surrogate of CaCo-2-formed 3D epithelial architecture. PHK26-labeled PBMC were manually counted on each imaging plane, and the ratio of the cells that were trapped in endothelium channel or have successfully infiltrated into the membrane and epithelium channel was quantified. For apparent permeability (P_app_) measurement, 50 µg/ml Cascade Blue Dextran 3000 MW (Thermo Fisher) was added in the CaCo-2 cell media and the flow-through from both channels including inlets and outlets was collected daily. The fluorescence intensity was measured on a Tecan microplate reader. The P_app_ values were calculated according to manufacturer’s protocol to track the development of a healthy membrane barrier as well as to evaluate the severity of barrier function disruption.

### Mixed Lymphocyte Reaction

Human peripheral blood mononuclear cells were isolated from Buffy Coats from anonymous donors (DRK-Blutspendedienst Baden-Württemberg-Hessen, Institut für Transfusionsmedizin und Immunhämatologie, Frankfurt, Germany) using Ficoll-Paque density gradient centrifugation. Isolated PBMCs from 2 non-related donors were co-cultured in pairs at a concentration of 2 x 10^6^ cells/ml in 48 well plates (final volume 250 µl) for 6 days in T-cell medium (RPMI 1640, penicillin (100DU/mL), streptomycin (100 μg/mL), FCS (10%), non-essential and essential amino acids (1%), sodium pyruvate (1%) and 1% 4-(2-hydroxyethyl) -1 piperazineethanesulfonic acid (HEPES)). rhIL-2 (10 ng/mL; PrepoTech) and β-mercaptoethanol (50 μM; Gibco) were added at day 0, 2, and 4. Cells were treated daily with full-length IL-38^19^, human recombinant IL-1RA (Peprotech), or human recombinant IL-36RA (Peprotech) at concentration 100 ng/ml. Supernatants were collected every day for cytokine measurements starting on day 1. Cells were harvested daily for gene expression analysis or were analyzed on day 6 by flow cytometry.

### T-cell isolation, activation and treatment

Primary human peripheral blood cells were isolated from Buffy Coats of anonymous donors (DRK-Blutspendedienst Baden-Württemberg-Hessen, Institut für Transfusionsmedizin und Immunhämatologie, Frankfurt am Main). T-cells were isolated using the EasySep™ Human T-cell Isolation Kit (Stemcell Technologies) through negative selection. The purity of T-cells was greater than 95%, as confirmed by flow cytometry. The cells were cultured at concentration 1 × 10^6^ cells/mL in T-cell medium. Cells were supplemented with human recombinant IL-2 and β-mercaptoethanol at days 0, 2, and 4. Cells were cultured for up to 6 days and treated daily with +/- IL-38 (100 ng/ml). Supernatants were collected on days 2, 4 and 6 for cytokine measurement.

### Monocyte isolation, activation

Primary human peripheral blood cells were isolated from Buffy Coats of anonymous donors (DRK-Blutspendedienst Baden-Württemberg-Hessen, Institut für Transfusionsmedizin und Immunhämatologie, Frankfurt am Main). Monocytes were isolated via human CD14 Microbeads (Miltenyi Biotec) and cultured in RPMI 1640 media (penicillin (100 U/mL), streptomycin (100 μg/mL), FCS (10%)) supplemented with rhM-CSF (50 ng/ml, Immunotools) for 24 hours at concentration 1 × 10^6^ cells/mL. Cells were pretreated with IL-38 (100 ng/ml, Adipogen) before stimuli (LPS (10 ng/ml) and TNF-α (10 ng/ml, Immunotools) were added. After an overnight culture, cells were collected and analyzed by flow cytometry.

### Macrophage differentiation

Human peripheral blood mononuclear cells were isolated from buffy coats of anonymous donors (DRK-Blutspendedienst Baden-Württemberg-Hessen, Institut für Transfusionsmedizin und Immunhämatologie, Frankfurt, Germany) using Ficoll density gradient centrifugation. Peripheral blood mononuclear cells were washed twice with PBS containing 2 mM EDTA and thereafter incubated for 2 h under growth conditions in RPMI 1640 (penicillin (100 U/mL), streptomycin (100 μg/mL)) to enable adherence to culture dishes (Sarstedt, Nümbrecht, Germany). Non-adherent cells were removed. Monocytes were differentiated into naïve macrophages with RPMI 1640 media containing 3% AB-positive human serum (DRK-Blutspendedienst Baden-Württemberg-Hessen, Frankfurt, Germany) for up to 7 days. Alternatively, macrophages were differentiated from monocytes isolated with human CD14 Microbeads (Miltenyi Biotec). Cells were cultured in RPMI 1640 media (penicillin (100 U/mL), streptomycin (100 μg/mL), FCS (10%)) supplemented with recombinant human M-CSF (50 ng/ml, Immunotools) for 3 and 7 days. The media was changed on days 2 and 5 with fresh recombinant human M-CSF. Treatments were added daily. Cells in both protocols were harvested on days 3 and 7 and inspected by gene expression analysis and by flow cytometry.

### Antigen-presentation assay

Primary human peripheral blood cells were isolated from Buffy Coats of anonymous donors (DRK-Blutspendedienst Baden-Württemberg-Hessen, Institut für Transfusionsmedizin und Immunhämatologie, Frankfurt am Main). T-cells were isolated using the EasySep™ Human T-cell Isolation Kit (Stemcell Technologies) through negative selection. T-cells were cultured alone for 3 days in the T-cell medium supplemented with rhIL-2 (10 ng/ml, Peprotech), rhIL-7 (10 ng/ml, Peprotech) and β-mercaptoethanol. Macrophages were differentiated from primary human monocytes as indicated above. On day 2, CD14+ cells were pulsed with Tetanus Toxoid (1 µg/ml, Sigma-Aldrich) and PepTivator® CMV pp65 (1 µg/ml, Miltenyi Biotec) for 24 h. Afterwards, cells were washed 2 times with PBS. T-cells were stained with cell proliferation dye eFluor 670 according to the manufacturer’s description and seeded on top of early macrophages at a ratio of 1:2 (macrophages: T-cells) for 3 days. IL-38 was added daily to CD14+ cells at a concentration of 100 ng/ml. T-cell proliferation was analyzed by flow cytometry.

### Cytometric Bead Array

Human GrzB, IFN-γ, TNF-α, IL-6, IL-10, IL-17A concentrations in MLR, T-cell- and Organ-on-Chip-derived supernatants and murine IFN-γ, TNF-α, IL-6 IL-17a, IL-10, MCP-1 and IL-1β in mouse serum were measured using Cytometric Bead Array Flex Sets (BD Bioscience) via flow cytometry. Data were analyzed using FlowJo V.10 (BD Biosciences).

### Flow Cytometry

Human cells were harvested, pelleted by centrifugation, blocked with FcR-blocking reagent (Miltenyi Biotec) in 0.5% PBS-BSA, stained with fluorochrome-conjugated antibodies and analyzed on a FACSymphony A5 flow cytometer (BD Biosciences). For intracellular FoxP3 staining, cells were fixed and permeabilized with Cytofix/Cytoperm (BD) according to manufacturer’s instruction and subsequently blocked with FcR-blocking reagent and stained with anti-FoxP3 antibody. For intracellular cytokine staining, single-cell suspensions were treated with 5 μg/mL brefeldin A (eBioscience), Golgi stop and PMA/ionomicyn (BD Biosciences) for 4 hours at 37°C followed by cell surface marker staining. Then, cells were fixed, permeabilized with Cytofix/Cytoperm (BD), and stained with anti-IFN-γ, anti- TNF-α, anti-IL-10, and anti-IL-6 antibodies. Gastrointestinal (GI) murine samples were processed using Lamina-Propria Dissociation Kits (Miltenyi Biotec) and the gentleMACS Dissociator (Miltenyi Biotec) according to the manufacturer’s protocol. Single cell suspensions from processed GI, blood, spleen, and liver samples were filtered through 70 μm cell strainers (BD Biosciences), blocked with 2% murine Fc receptor binding inhibitor (Miltenyi Biotec) and stained with antibodies (Suppl. Table 2). Compensation Beads (BD) were used for single color compensation to create multicolor compensation matrices. For gating, fluorescence minus one controls were used.

### Quantitative Real-time PCR

Cells were lysed 400 µl TRIzol (Invitrogen) and RNA was isolated as per manufacturer recommendations. mRNA (1 µg) was used to perfrom reverse transcription with the Maxima First Strand cDNA Synthesis kit (Thermo Fisher Scientific). Quantitative real-time PCR reactions were done with the SYBR Select Master Mix, and the QuantStudio 5 Real-Time PCR System (Thermo Fisher Scientific). Relative mRNA expression was analyzed based on the ΔΔcycle threshold method and normalized to *TBP* as housekeeping gene.). The following human primers were used:

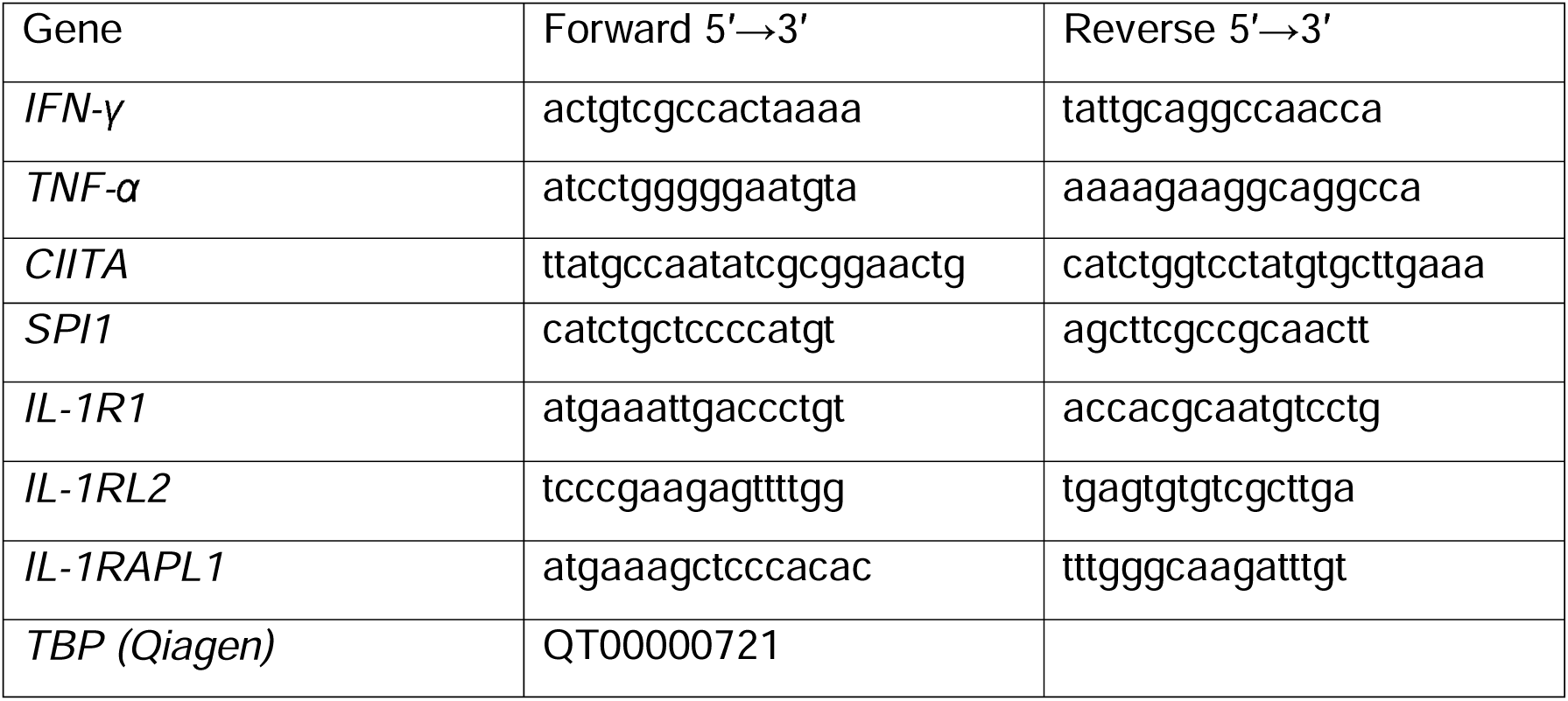

### Immunocytochemistry and confocal microscopy

Macrophages were pretreated with 10 µg/ml of IL-38 for 1 hour. After, cells were fixed in 4% paraformaldehyde and permeabilized with 0.1% triton X for 10 min (except for non-permeabilized controls), blocked with 5% BSA containing 100 mM Glycine and human FcR Blocking Reagent (1:50 dilution, Miltenyi Biotec). Subsequently, cells were incubated with 1:100 anti-Hu-IL-38 antibody (Invitrogen) at 4 °C overnight in PBS containing 1% BSA, FcR Blocking Reagent and 0,05% triton X. The following day cells were counterstained with 20 nM MEMGlow 488 (Cytoskeleton) for 10 min and 1 µg/ml DAPI for 1 min. Images were acquired using Zeiss LSM 800 Axio-Observer with Plan-Apochromat 40×/1.4 Oil DIC M27 objective and GaAsP detector. Acquisition and airyscan processing was performed using Zen Blue software. Images were processed with ImageJ.

### Immunohistochemistry

For liver and GI tissue sections, the following murine primary antibodies were used: CD3 (Abcam), CD8 (Cell signaling), CD4 (Cell Signaling), FoxP3 (Cell Signaling), MHC-II (Invitrogen), and F4/80 (Cell Signaling). Detailed information regarding the antibodies used is provided in Supplemental Table 3. The PhenoImager HT automated quantitative pathology imaging system (Akoya) was used for image acquisition at ×20 and ×40, and images were analyzed using inForm 2.6 Software (Akoya Bioscience).

### Cellular Indexing of Transcriptomes and Epitopes: library preparation and sequencing

PBMCs from anonymous donors were isolated via Ficoll gradient centrifugation and maintained according to standard procedure. The cells were co-cultured in pairs for 6 days to identify the strongest allogeneic response by measuring IFN -γ and TNF-α levels in the supernatants. After 4 combinations were chosen, cells were thawed and co-cultured to collect samples for CITE-seq. Single-cell suspensions were harvested on days 0, 3, and 5 and purified with the Dead Cell Removal Kit (Miltenyi). After dead cell removal, cells were labelled with 10X Genomics CMOs (Day 0) and antybodies (BioLegend TotalSeq B Human Universal Cocktail V1), or with antibodies only (day 3 and day 5), and processed on 10x Genomics according to manufacturer’s instruction. Single-cell capture and library preparation were performed using the Chromium Single Cell 3′ v3.1 kit (10x Genomics) following the manufacturer’s instructions. Libraries were sequenced on an Illumina NextSeq 2000 platform using the p3 reagent kit. Raw FASQ files were processed using Cell Ranger, and cell calling matrices were generated using Cell Ranger as well.

### CITE-sequencing analysis

Downstream analyses were conducted using Seurat (v.4)^36^ pipeline. The datasets were quality-controlled via filtering out cells containing RNA feature < 200 and mitochondrial RNA < 10%. RNA data were normalized using SCTransformation v.2, retaining the top 2000 variable genes. ADT (Antibody-derived tags) were normalized using the CLR function. Data integration was applied within the Seurat workflow. The batch correction and doublet control were performed using packages CellMixS^37^ and Doubletfinder^38^. Unsupervised clustering was based on the top 15 principal components, and UMAP was used for dimensionality reduction and visualization. Integrated in the Seurat FindAllMarkers function and Clustree^39^ were applied to identify and annotate major cell populations presented in our dataset. Differential gene expression analysis was performed using the implemented Seurat workflow function FindMarkers. GO (gene ontology) and GSEA (gene set enrichment analysis) were scored based on significant differential gene expression (log2FC > 0.5, and p.value < 0.05). Inflammation and T-cell signatures were calculated with the AddModuleScore function in Seurat. Average expression was calculated using the AverageExpression function in Seurat.

### Statistical analysis

Data are presented as mean ± SEM. Statistically significant differences between the two groups were calculated using paired or unpaired two-tailed Student’s t-test. A one-sample t-test was used to calculate differences in normalized data. For multiple comparisons, multiple t-tests, one-way and two-way analysis of variance were used, followed by appropriate post-correction analysis. Statistical survival analysis was determined via log-rank test and correlation using the Spearman test. Statistical analysis was performed with GraphPad Prism V.9 and p values < 0.05 were considered statistically significant.

## Discussion

Acute GvHD remains a severe jeopardizing complication after allo-HST leading to high mortality. Despite the current advanced prophylaxis up to 60% of patients are at risk of developing GvHD^40^, indicating a need for new prophylactic and therapeutic strategies. The role of IL-38 in GvHD was previously unreported. Interfering with other IL-1 cytokine family members in the context of GvHD has yielded controversial results. No therapeutic effect of IL-1RA was observed in patients undergoing allo-HST, and approximately half of them developed severe disease^41^. This is in contrast to data in mouse models showing that blockade of IL-1 receptor attenuated disease score in animals via inhibiting Th17 generation and expansion of regulatory T-cells^42^. Moreover, depletion of IL-1β or receptor antagonism ameliorated GvHD-associated mortality in mouse models^43^. Our study provides evidence for a prophylactic potential of IL-38 in GvHD, associated with increased Treg formation and an alleviated disease score. Regulatory T-cells have been shown to prevent and ameliorate acute GvHD in murine models, and elevated proportions of Tregs were correlated with a better outcome in patients after allo-HSCT^44–47^. Adoptive Tregs transfer has emerged as a promising strategy for the prevention and treatment of GvHD^48^. Due to low cell numbers, several protocols of Treg expansion were successfully established. However, this strategy comes at a considerable cost, and in vitro expanded Tregs are prone to having unstable FoxP3, which limits immunosuppressive function of Tregs in the context of GvHD^49^. Thus, there is a clear unmet need for novel compounds that drive Treg expansion *in vivo*. IL-38 was able to expand Tregs at the systemic level, as well as in the periphery (GI and liver), which correlated with the amelioration of the disease in IL-38-treated mice. Moreover, we observed an increase in CD8+ regulatory T-cells (CD8+FoxP3+) in LPL of IL-38-treated mice, which were previously shown to improve survival while preserving GvL activity in a murine model of GvHD^50^. Our findings are partly supported by previous studies. IL-38 overexpressing DCs were able to expand Tregs and ameliorate asthma in a murine model^51^, where IL-38 had a direct effect on Tregs isolated from murine splenocytes *in vitro*, inducing transcription of FoxP3 and CTLA-4, and increasing secretion of IL-10 and TGF-β1^51^. In line with these findings, our data show that IL-38 is prone to induce Tregs, also in an allogeneic MLR, albeit at a minor level, which may be due to the limited time-course of these assays.

Besides generation of Tregs, modulation of antigen-presentation activity is one of the attractive approaches in prophylaxis and treatment of GvHD. Pro-inflammatory macrophages are predominant in acute GvHD, whereas anti-inflammatory macrophages expand during chronic disease^52^. Depletion of macrophages^53^ showed worsened survival of mice in humanized model of GvHD. Thus, alternative strategies are needed to target APCs. Here, we show that IL-38 transiently alters MHC-II expression on early and differentiated macrophages in the human *in vitro* setting. Our data showing IL-38 alters MHC-II in myeloid cells is partly confirmed by another study. Here, IL-38 demonstrated to reduce antigen presentation capacity in PBMCs and decreased the expression of surface co-stimulatory molecules in monocyte-derived DCs^54^. The mechanism underlying this modulation is still unclear. IL-38 has been postulated as a partial antagonist to IL-36R and IL-1R since it shares 40% of homology with IL-36RA and IL-1RA cytokines. IL-38 showed a similar dose-response effect to IL-36RA in *C. albicans*-stimulated human PBMCs^24^. However, this notion is challenged by recent evidence demonstrating that IL-38 failed to antagonize cytokine production through IL-36R and IL-1R1 signaling in IL-36R- and IL-1R1-expressing cell lines^25^. In our *in vitro* data, only IL-38, but not IL-1RA and IL-36RA, attenuated inflammatory responses in MLR. Moreover, HLA-DR expression on the surface of macrophages was explicitly reduced only in IL-38-treated cells. Thus, IL-38 appears to employ unique signaling pathways independent of IL-1 and IL-36 receptors. IL-38 has previously shown intracellular activity, as suggested by our data as well^60–61^. Our findings are partially supported by previous studies demonstrating that IL-38 modulates trained immunity in macrophages through epigenetic regulatory mechanisms^57^. Importantly, IL-38 affected antigen-presentation capacity at early time-points of macrophage differentiation. Since we analyzed at a late stage of GvHD progression in the mouse model, major effects on GFP+ T-cells were not observed. Similarly, we observed only a moderate increase in Tregs on day 5 in human MLRs, indicating that the impact of IL-38 on Treg formation occurs at a later stage. Moreover, Treg expansion requires a complex tissue microenvironment, which is found in lymphoid organs rather than *in vitro* assays with reduced complexity^49,58,59^. Further studies are needed to investigate the mechanisms underlying the immunosuppressive action of IL-38 in GvHD. Previously, IL-38 was shown to diminish inflammation in DSS-induced colitis in mice^60^ and to regulate homeostasis in intestinal stem cells^61^, which particularly highlights the importance of IL-38 to modulate GI-GvHD. Given that IL-38 expression is elevated in differentiated keratinocytes of healthy skin compared to lesional skin in cutaneous GvHD, exogenous IL-38 administration may represent a promising therapeutic strategy to attenuate inflammation in this condition. IL-1RAPL1 (IL-1R9) is one of the potential IL-38 receptors, predominantly expressed in the central nervous system (CNS)^62^, and is associated with autism spectrum disorder (ASD)^63^. IL-38 diminished microglial inflammatory mediators^64^. These properties make IL-38 an attractive therapeutic candidate in the context of CNS GvHD^65^.

Overall, we demonstrate IL-38 as a prophylactic compound in the context of murine GvHD and in human GvHD models, employing a unique signaling pathway. The relative contribution of altered antigen presentation versus expansion of Tregs remains to be determined, but IL-38 treatment may represent an attractive approach against acute GvHD, also in combination with other current strategies.

## Acknowledgments

The authors thank Andrew Vorobyov and Margarethe Mijatovic for excellent technical assistance.

## Author Contributions

Conceptualization, A.K., W.X., K.I., and A.W.; methodology, A.K., W.X., I.M., N.B., D.N., I.K., B.A., K.I., A.W.,; formal analysis, A.K., W.X., I.M., K.I., and A.W.; investigation, A.K., W.X., K.I., MAF. E., AC. J., I.K., and B.A.; resources, I.M., K.I., B.B., and A.W.; data curation, A.K., W.X., I.M., K.I., and A.W.; writing—original draft preparation, A.K., and A.W.; writing—review and editing, all authors; visualization, A.K., K.I., and A.W.; supervision, B.B., and A.W.; funding acquisition, B.B., and A.W. All authors have read and agreed to the published version of the manuscript.

## Data Availability Statement

The transcriptomic datasets generated during and analyzed during the current study are available at GEO: GSE335061. All other datasets generated during and/or analyzed during the current study are available from the corresponding author upon reasonable request.

