## Supplementary Figures and Figure Legends, Supplementary Methods for "IL-38 limits alloreactivity through modulating myeloid and T-cell activation"

**Supplementary material**

**Supplementary Figures and Figure Legends**

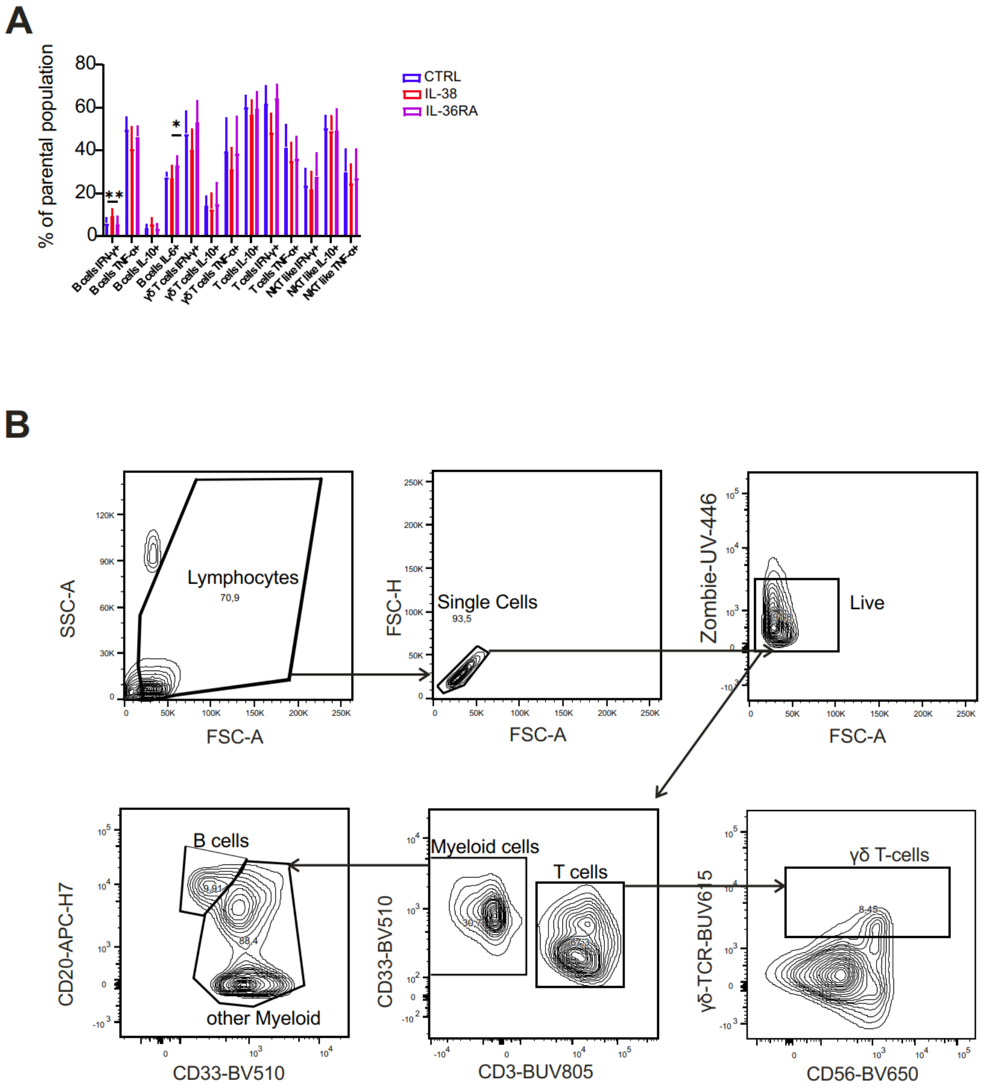


**Figure S1. Impact of IL-38 and IL-36Ra on immune cell populations in two-way MLR.** Human PBMCs from unrelated donors were co-cultured for 6 days and treated with IL-38 or IL-36RA (each at 100 ng/ml). Cells were collected on day 6 and analyzed by intracellular FACS. **(A)** The graph shows the percentage of main immune cell populations expressing intracellular-stained cytokines obtained in MLR and determined by flow cytometry. **(B)** Representative FACS plots of intracellular staining of MLR assay gating strategy.

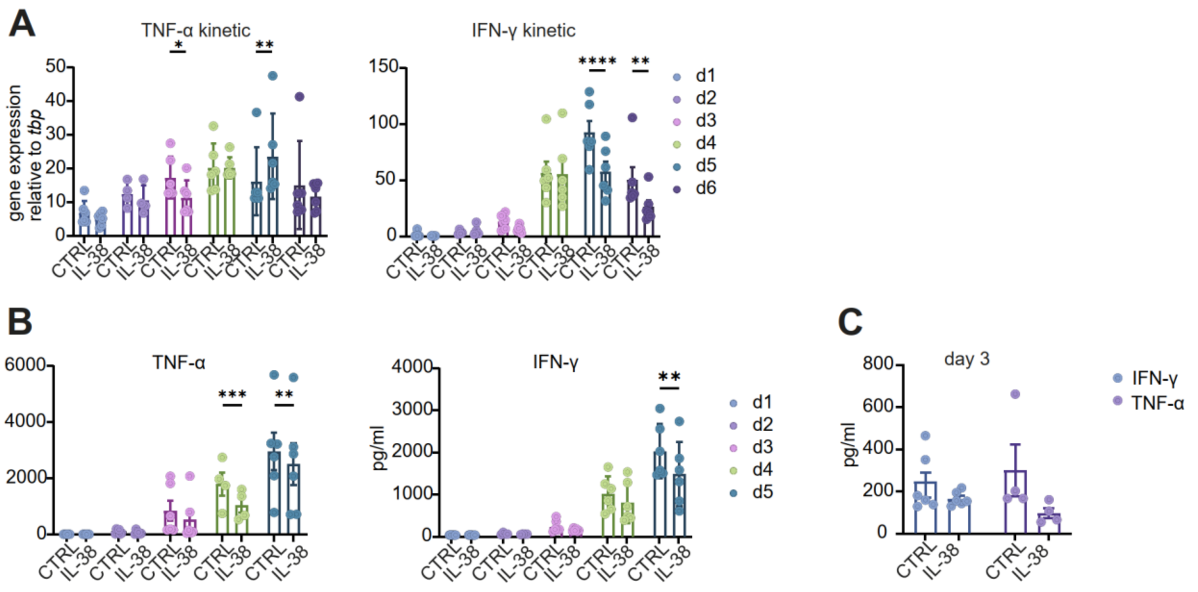


**Figure S2. Expression kinetics of IFN-γ and TNF-α in MLR in the presence of IL-38.** Human PBMCs of unrelated donors were co-cultured for 6 days and treated with IL-38 at a concentration of 100 ng/ml**.** mRNA was collected on days 1-6 during the MLR assay **(A).** TNF-α (left panel) and IFN-γ (right panel) mRNA expression were determined by qPCR. **(B)** Supernatants were collected on days 1-6 during the MLR assay**.** The levels of IFN-γ (left panel) and TNF-α (right panel) were determined by flow cytometry, presented as mean fluorescent intensity (MFI). **(C)** The graph shows the MFI of TNF-α and IFN-γ measured in the supernatants collected on day 3. The data are from one representative experiment out of three. Each data point corresponds to one independent donor pair (n = 6). Data are shown as mean ± SEM. **p* < 0.05, ***p* < 0.01, ****p* < 0.001, *****p* < 0.0001; *p*-values were calculated using two-way ANOVA.

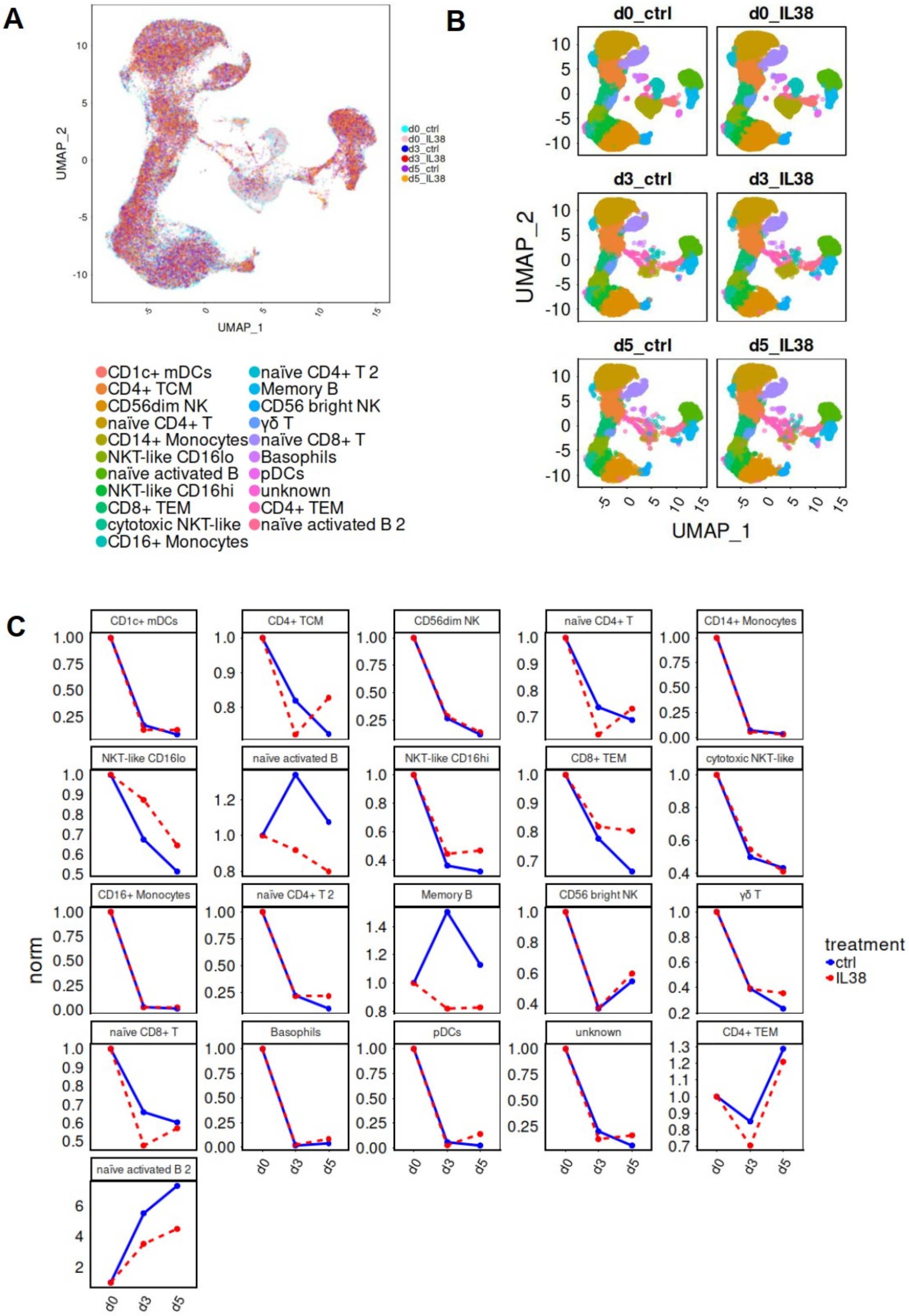


**Figure S3. Impact of IL-38 on immune cell abundance in human MLR.** Human PBMCs from 4 unrelated donors were co-cultured in pairs and treated with IL-38 (100 ng/ml) daily. Cells were collected on days 0, 3, and 5 and analyzed by CITE seq. **(A)** UMAP plot of integrated data across all conditions. **(B)** UMAP plots of the integrated data shown by time point and conditions (d0_ctrl, d0_IL38, d3_ctrl, d3_IL38, d5_ctrl, d5_IL38) with annotated clusters. **(C)** The graph depicts the cell abundance across all time points between the control and IL-38 groups. Data are normalized to the counts of day 0.

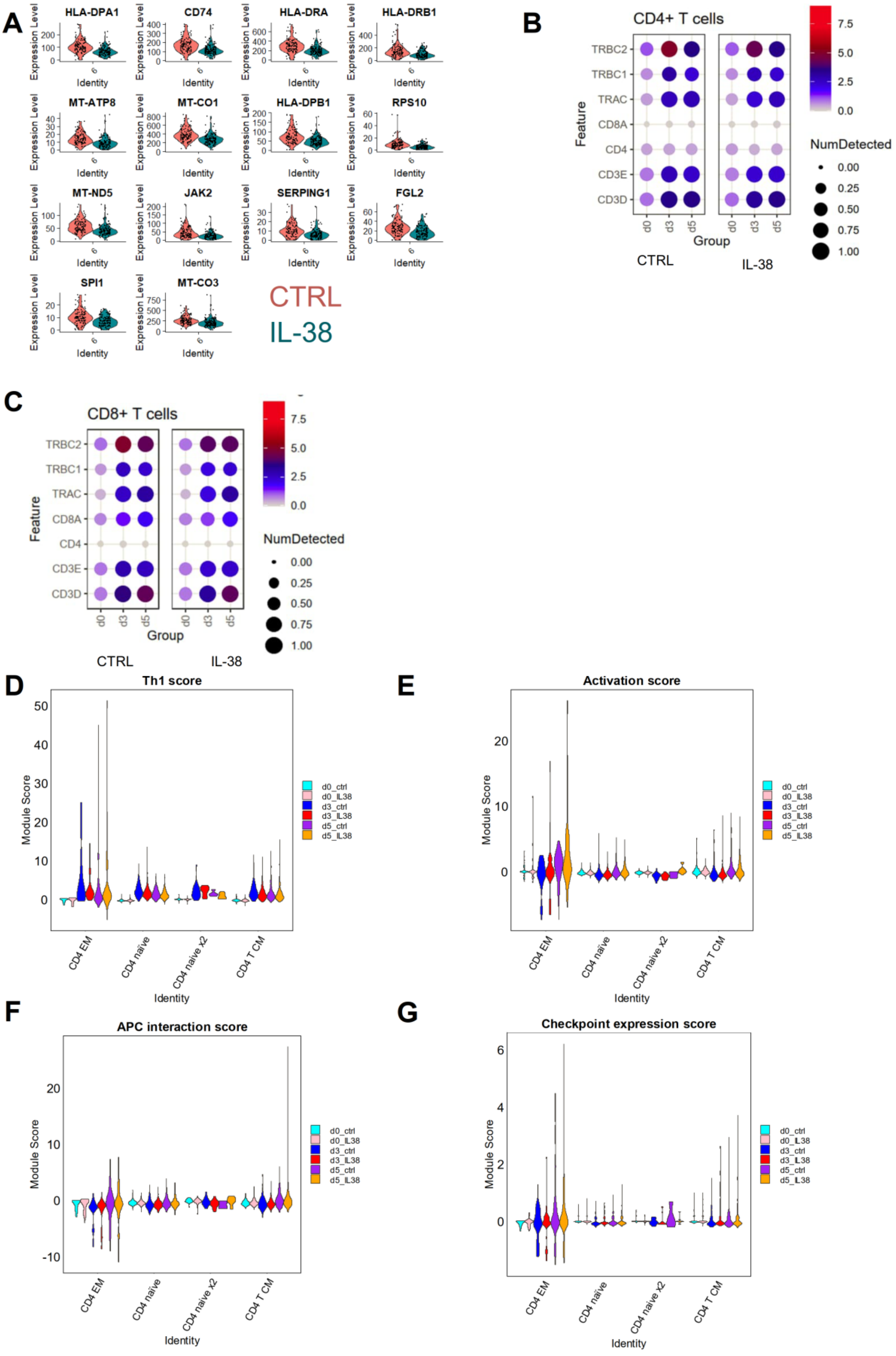
A
HLA-DPA1
CD74
HLA-DRA
HLA-DRB1
MT-ATP8
MT-CO1
HLA-DPB1
RPS10
MT-ND5
JAK2
SERPING1
FGL2
SPI1
MT-CO3
Expression Level
Identity
CTRL
IL-38
B
CD4+ T cells
TRBC2
TRBC1
TRAC
CD8A
CD4
CD3E
CD3D
Feature
d0
d3
d5
Group
CTRL
IL-38
7.5
5.0
2.5
0.0
NumDetected
0.00
0.25
0.50
0.75
1.00
C
CD8+ T cells
TRBC2
TRBC1
TRAC
CD8A
CD4
CD3E
CD3D
Feature
d0
d3
d5
Group
CTRL
IL-38
7.5
5.0
2.5
0.0
NumDetected
0.00
0.25
0.50
0.75
1.00
D
Th1 score
Module Score
CD4 EM
CD4 naïve
CD4 naïve x2
CD4 T CM
Identity
d0_ctrl
d0_IL38
d3_ctrl
d3_IL38
d5_ctrl
d5_IL38
E
Activation score
Module Score
CD4 EM
CD4 naïve
CD4 naïve x2
CD4 T CM
Identity
d0_ctrl
d0_IL38
d3_ctrl
d3_IL38
d5_ctrl
d5_IL38
F
APC interaction score
Module Score
CD4 EM
CD4 naïve
CD4 naïve x2
CD4 T CM
Identity
d0_ctrl
d0_IL38
d3_ctrl
d3_IL38
d5_ctrl
d5_IL38
G
Checkpoint expression score
Module Score
CD4 EM
CD4 naïve
CD4 naïve x2
CD4 T CM
Identity
d0_ctrl
d0_IL38
d3_ctrl
d3_IL38
d5_ctrl
d5_IL38

**Figure S4. Impact of IL-38 on myeloid and T-cell subsets.** Human PBMCs from 4 unrelated donors were co-cultured in pairs and treated daily with IL-38 (100 ng/ml). Cells were collected on days 0, 3, and 5 and analyzed by CITE seq.**(A)** Violin plots of down-regulated genes in CD14+ monocytes collected on day 3 (ctrl green, IL-38 pink). **(B)** The dot plot represents the expression of T-cell receptor, CD4, CD8, and CD3 genes of all pooled CD4+ T cell subpopulations. **(C)** The dot plot represents the expression of T-cell receptor, CD4, CD8, and CD3 genes of all pooled CD8+ T cell subpopulations. **(D-G)** The Th1 **(D)**, APC interaction **(E)**, Checkpoint expression **(F)**, and activation scores **(G)** were calculated across all CD4+ T-cell clusters (where d0_ctrl –cyan, d0_IL-38 – pink, d3_ctrl – blue, d3_IL-38 – red, d5_ctrl – violet, d5_IL-38 – yellow).

A

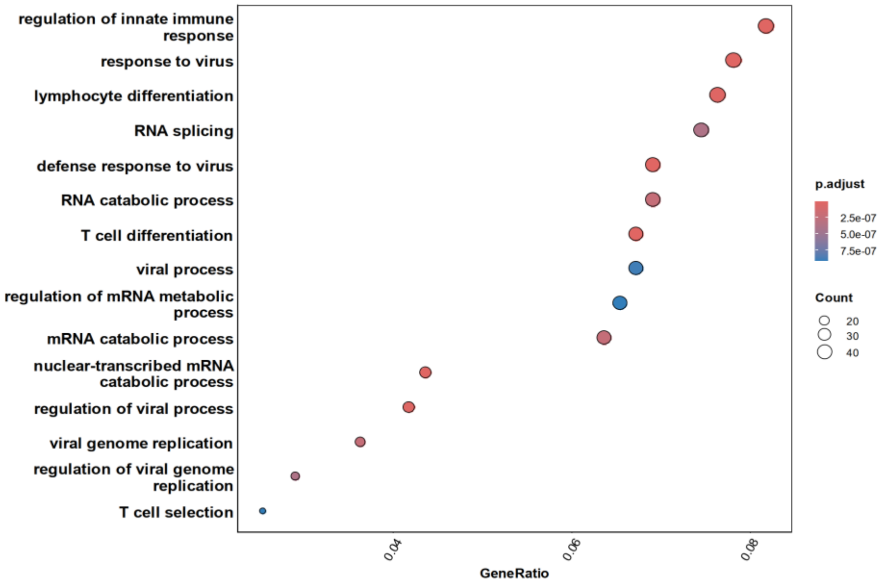


B

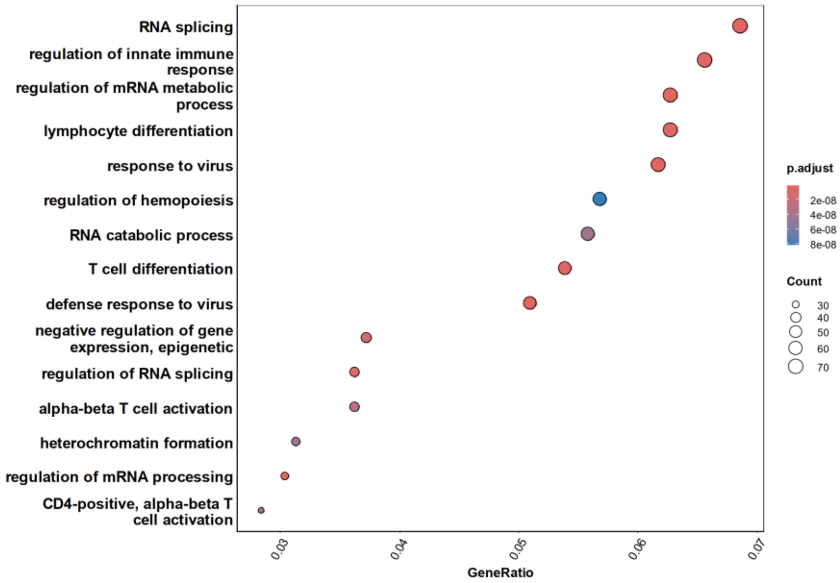


**Figure S5. IL-38 differently affects gene expression in CD4+ T-cell subsets on day 3 during MLR.** Human PBMCs from 4 unrelated donors were co-cultured in pairs and treated daily with IL-38 (100 ng/ml). Cells were collected on days 0, 3, and 5 and analyzed by CITE-seq.**(A)** Gene Ontology (GO) analysis of CD4+ T central memory cells collected on day 3 based on differential gene expression analysis between ctrl and IL-38. **(B)** Gene Ontology (GO) analysis of naïve CD4+ T-cells collected on day 3 based on differential gene expression analysis between ctrl and IL-38.

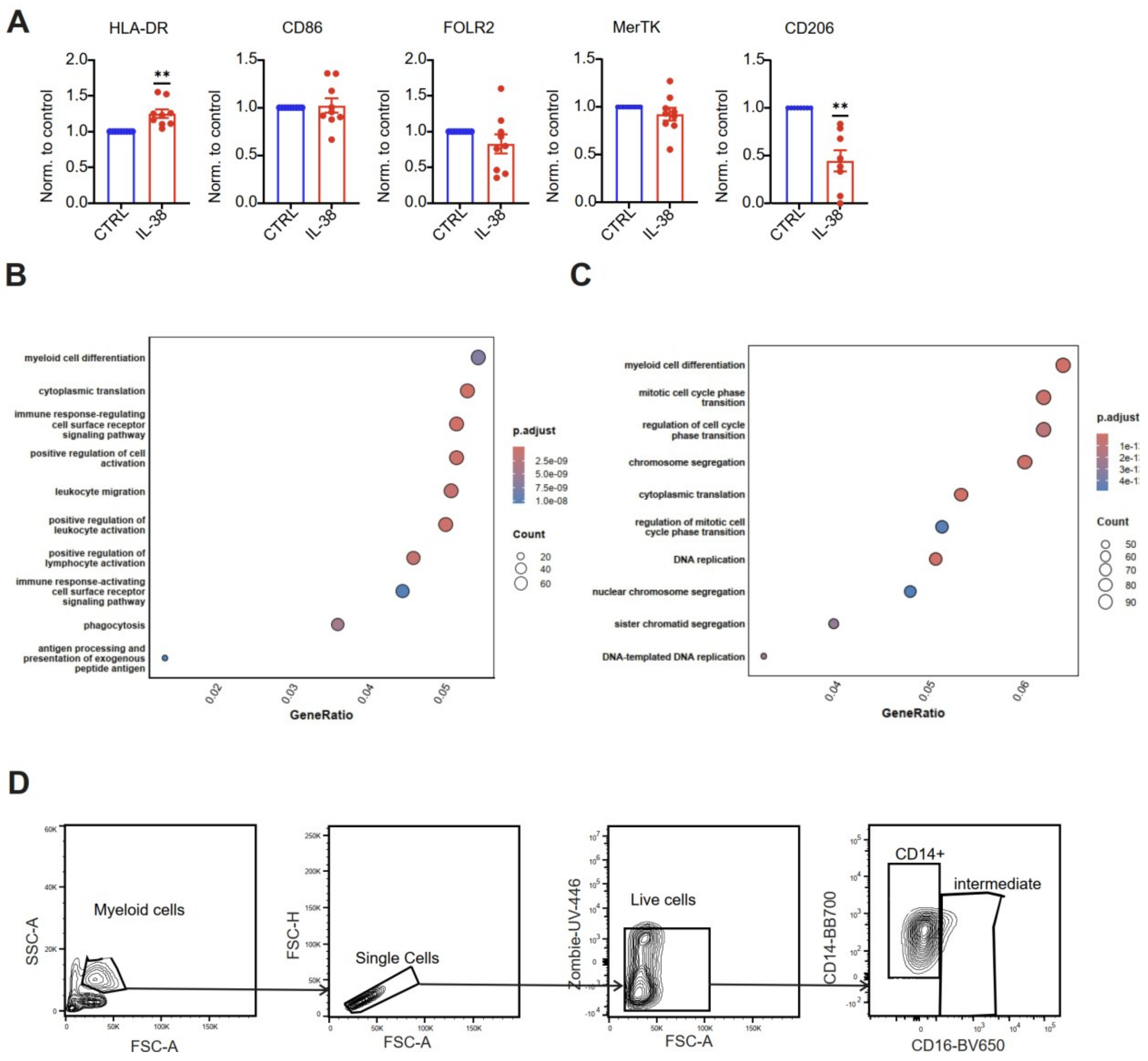


**Figure S6. IL-38 transiently modulates macrophage differentiation.** Monocytes were differentiated in the presence of human plasma up to 7 days and treated daily with IL-38 (100 ng/ml). **(A)** The surface expressions of HLA-DR, CD86, FOL2R, MerTK, and CD206 were determined by flow cytometry on day 7. Each data point represents an individual donor (n = 7), and the normalized data are from 3 independent experiments. Data are shown as mean ± SEM. *$p < 0.05$, **$p < 0.01$, ***$p < 0.001$, ****$p < 0.0001$; $p$-values were calculated using one-sample $t$-test. GO analysis of CD14+ monocytes based on differential gene expression analysis between day 3 and day 5 for IL-38 **(B)** and control **(C)** groups. **(D)** Representative FACS plots of macrophage differentiation assay gating strategy.

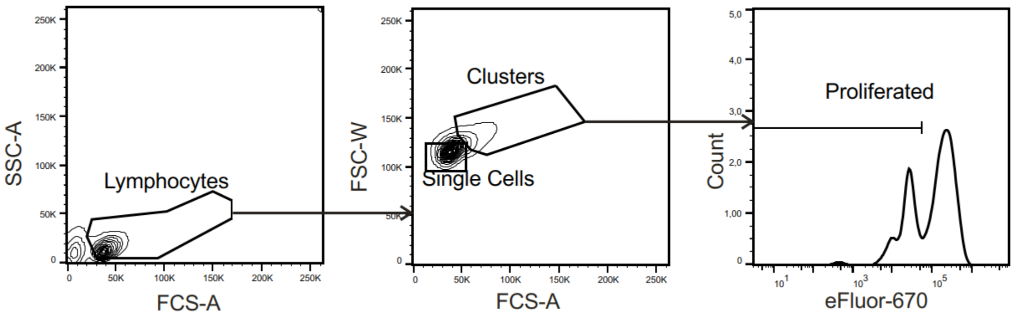


**Figure S7. Representative FACS plots of T-cell proliferation assay in one-way MLR experiments, gating strategy.**

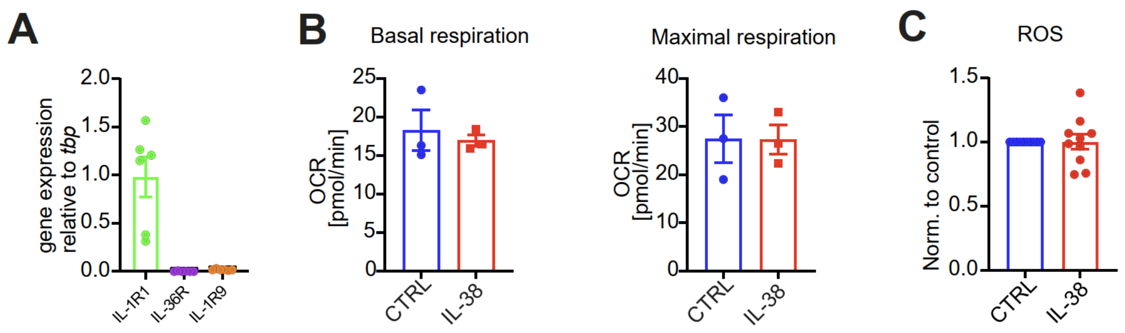


**Figure S8. Early macrophages do not show metabolic changes upon IL-38 treatment.** Monocytes were differentiated in the presence of human plasma for up to 3 days and treated daily with IL-38 (100 ng/ml). **(A)** Macrophages were collected on day 3 to investigate the gene expression levels of the main IL-38 receptors. mRNA expressions of IL-1R1, IL-36R and IL-1R9 were determined by qPCR. Each data point corresponds to one donor (n = 6). **(B)** The levels of basal and maximal respiration rates (OCR, oxygen consumption rate) in early macrophages were evaluated. IL-38 (100 ng/ml) was added daily. Each data point corresponds to one donor (n = 3). **(C)** Macrophages were differentiated in the presence of rhM-CSF (50 ng/ml) for up to 3 days in the presence of IL-38 (100 ng/ml), and cells were harvested to determine levels of reactive oxygen species (ROS) with CellROX Orange by flow cytometry.

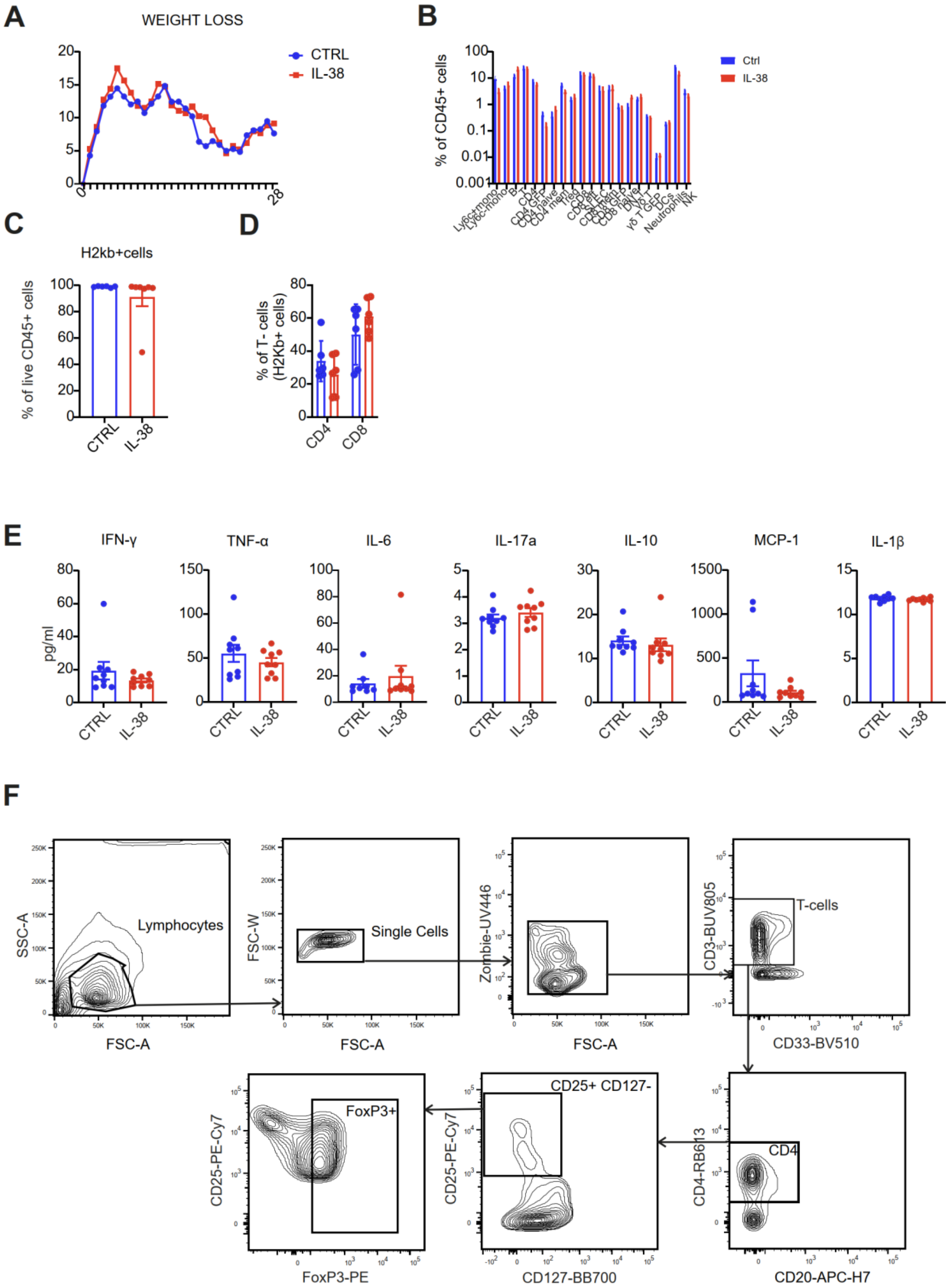


**Figure S9. Impact of IL-38 on murine GvHD.** BALB/c mice (host) were irradiated on day 0 and received T-cell-depleted bone marrow (BM) cells from wild-type (wt) C57BL/6 and T-cells from Nur77$^{GFP}$ mice on the following days. IL-38 was given (i.p) three times a week. **(A)** The graph shows the weight loss of animals over time between the control and IL-38 groups (n=9). **(B)** The immunofluorescent profiling of the immune donor-derived (H2kb+) cells in the blood between the control and IL-38 groups (n = 6). **(C)** The proportions of donor-derived immune cells of the total CD45+ cells in the blood (n = 6-7). **(D)** Percentages of CD4 and CD8 T-cells of donor-derived T-cells obtained from blood (n = 6). **(E)** Serum was analyzed via flow cytometry to evaluate concentrations of IFN-γ, TNF-α, IL-6, IL-17a, IL-10, MCP-1, and IL-1β (n = 9). Human PBMCs of unrelated donors were co-cultured for 6 days and treated with IL-38 at a concentration of 100 ng/ml. Cells were collected on days 3, 5, and 7 to validate regulatory T-cell formation. **(F)** Representative FACS plots of the human two-way MLR gating strategy. Each data point corresponds to an individual animal, and the data are from 3 independent experiments. Data are shown as mean ± SEM. *$p < 0.05$, **$p < 0.01$, ***$p < 0.001$, ****$p < 0.0001$; $p$-values were calculated using multiple $t$-test (B) and unpaired $t$-test (C-E).

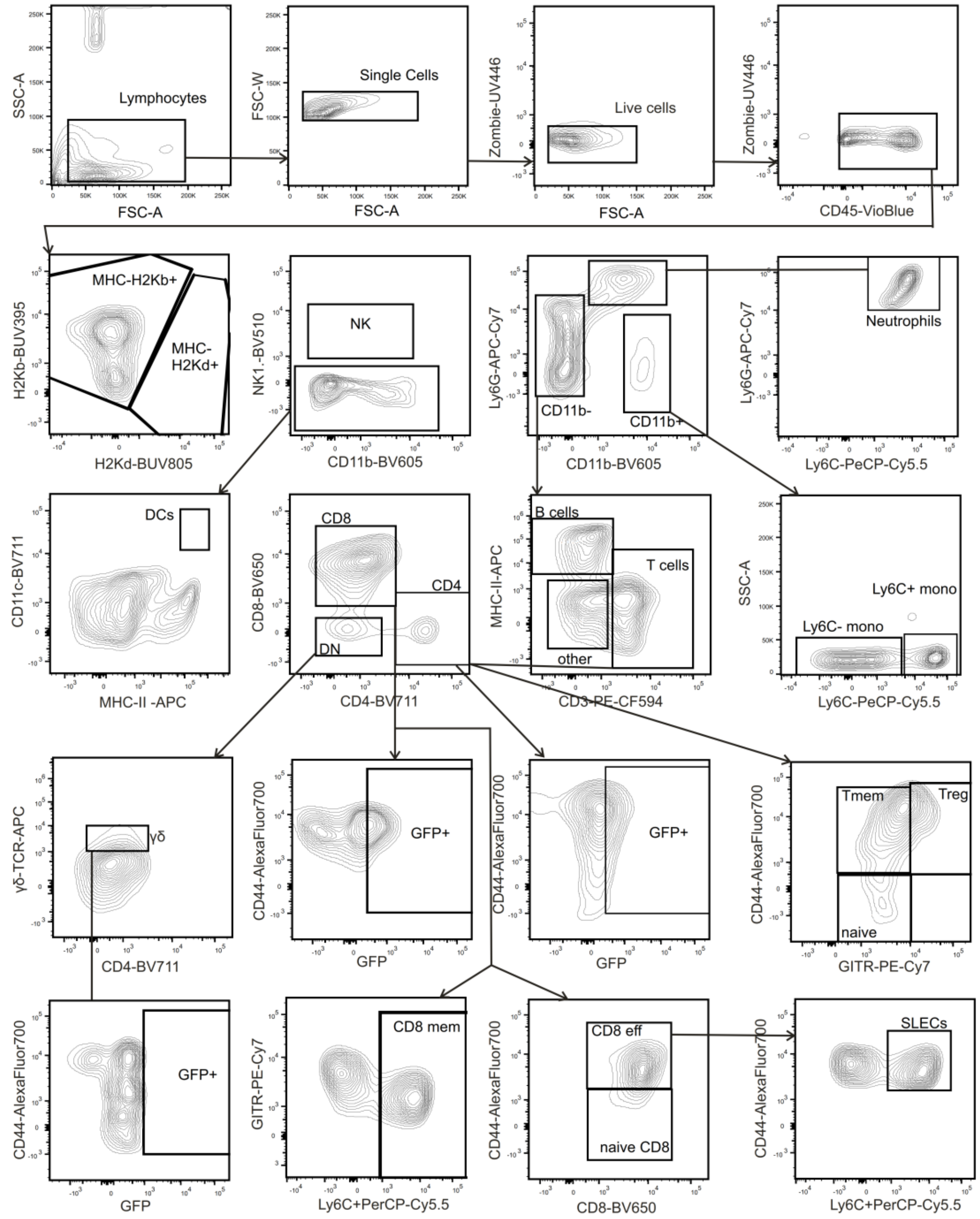


**Figure S10. Representative FACS plots of flow cytometric immunophenotyping of blood cells from murine GvHD samples.**

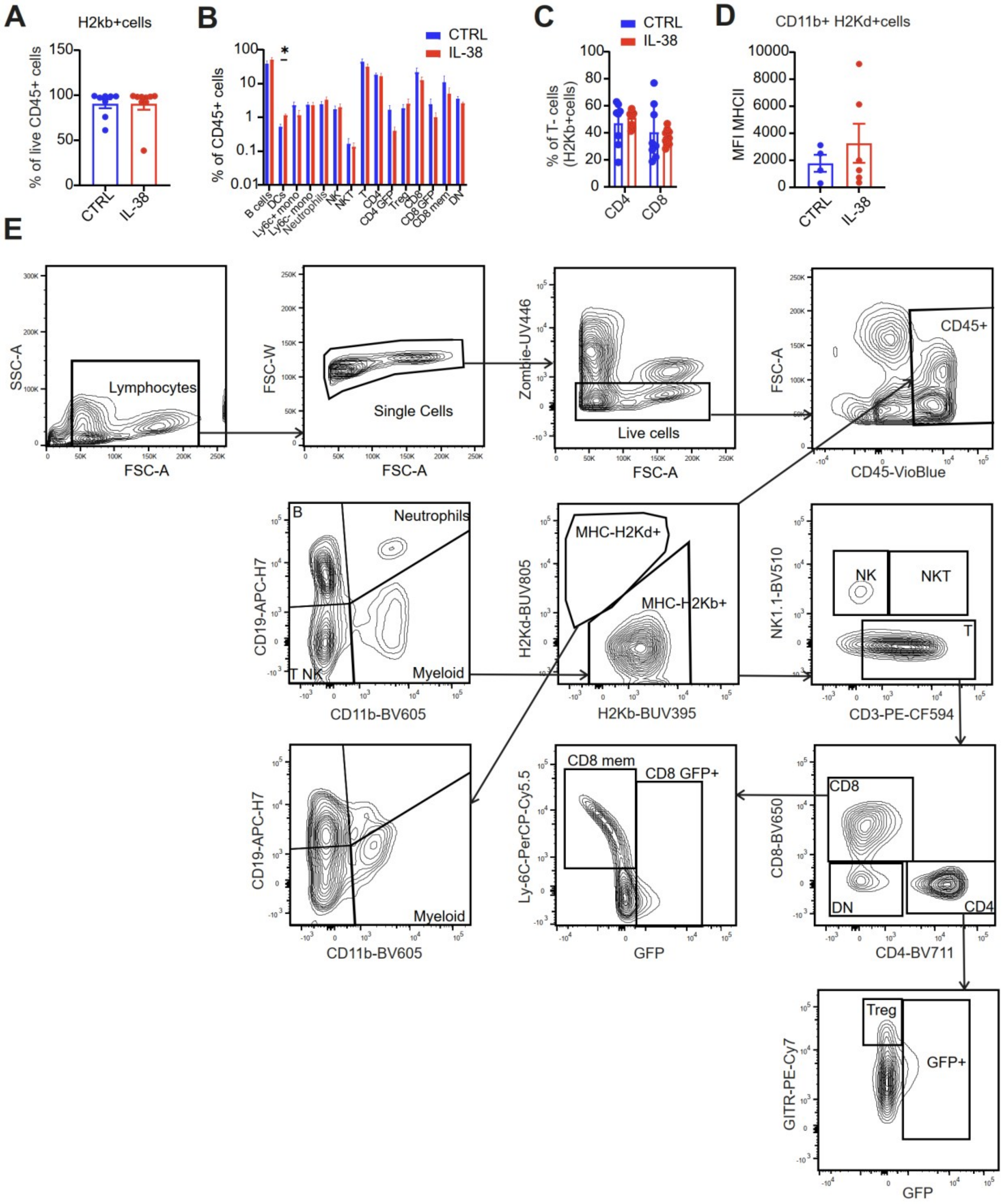


**Figure S11. Impact of IL-38 on immune cells in the spleen in GvHD.** BALB/c mice (host) were irradiated on day 0 and received T-cell-depleted bone marrow (BM) cells from wild-type (wt) C57BL/6 and T-cells from Nur77$^{GFP}$ mice on the following days. IL-38 was given (i.p) three times a week. **(A)** Proportions of donor-derived immune cells of the total CD45+ cells in the spleen determined by flow cytometry (n = 8-9). **(B)** Profile of donor-derived (H2kb+) immune cells in the spleen between the control and IL-38 groups (n = 8-9). **(C)** Percentages of CD4+ T-cells and CD8+ T-cells of donor-derived T-cells obtained from the spleen (n = 8-9) between the control and IL-38 groups. **(D)** The expression of MHC-II on the surface of host-derived APCs between control and IL-38 groups obtained from the spleen was determined by flow cytometry (n = 4-6). **(E)** Representative FACS plots of flow cytometric immunophenotyping of spleen cells from murine GvHD samples. Each data point corresponds to an individual animal, and the data are from 3 independent experiments. Data are shown as mean ± SEM. *$p < 0.05$, **$p < 0.01$, ***$p < 0.001$, ****$p < 0.0001$; *p*-values were calculated using multiple *t*-test (B) and unpaired *t*-test (A, C, D).

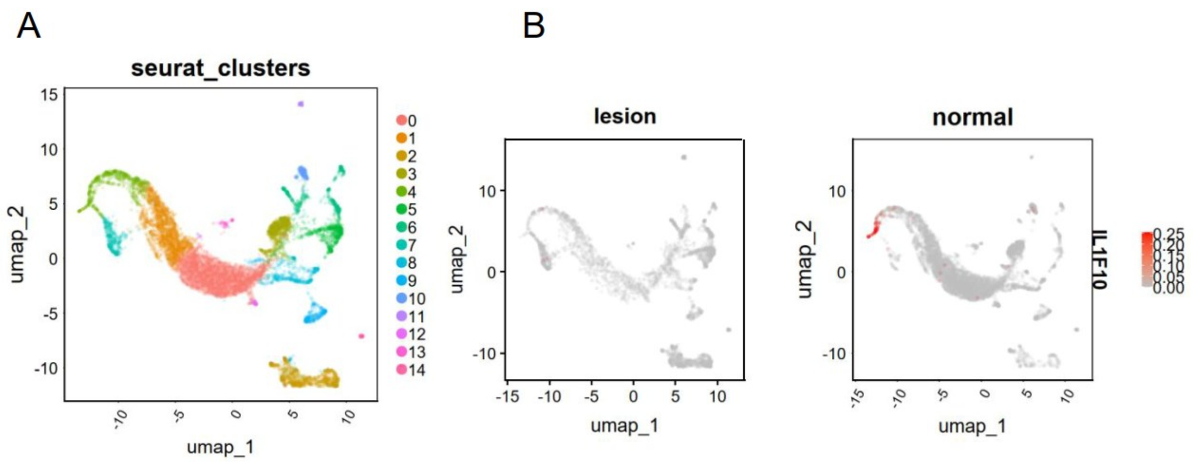


**Figure S12. IL-38 is expressed in the skin of healthy individuals compared to patients with lesional cutaneous GvHD.**

The single-cell transcriptomic data of epithelial samples from healthy donors and patients with lesional cutaneous GvHD were analyzed with the Seurat package. The datasets[1] were filtered and normalized. The FindAllMarkers function was used to identify major cell populations. The expression of IL-38 was validated between healthy and GvHD controls. **(A)** UMAP plot of integrated data of epithelial cells across healthy and lesional conditions. Cluster 3 is differentiated keratinocytes. **(B)** The feature plots of IL-38 (IL1F10) expression between healthy and lesional GvHD samples.

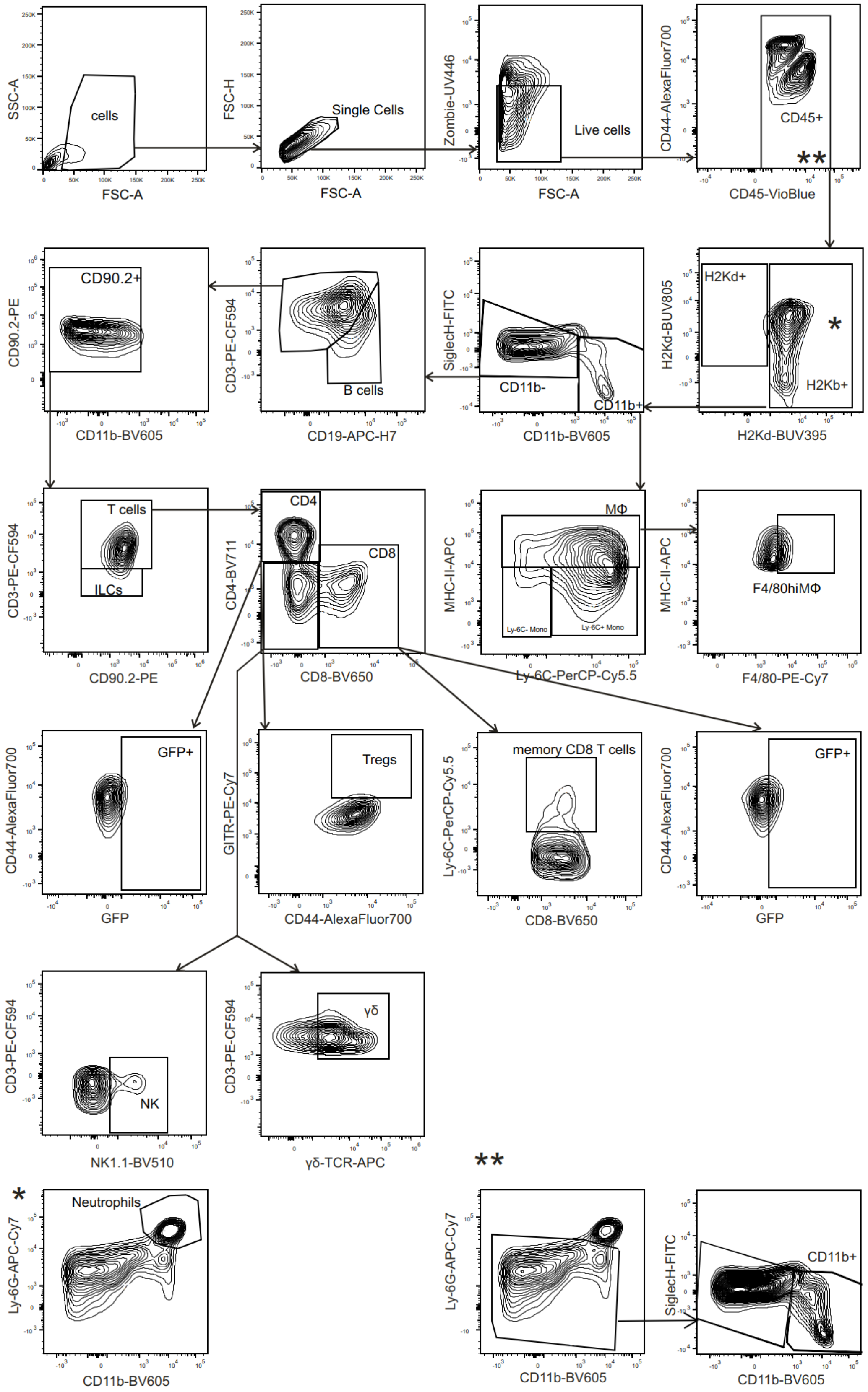


**Figure S13. Representative FACS plots of flow cytometric immunophenotyping of liver cells from murine GvHD samples.**

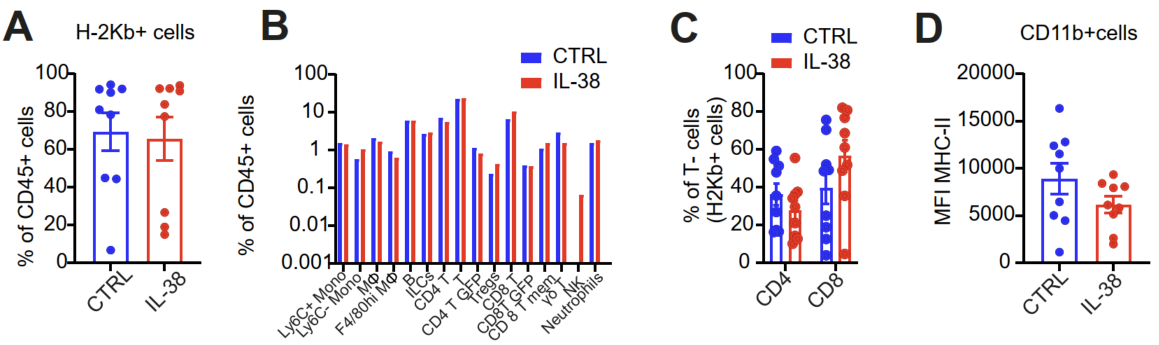


**Figure S14. IL-38 affects immune cells and host-derived cells in the liver during GvHD.** BALB/c mice (host) were irradiated on day 0 and received T-cell-depleted bone marrow (BM) cells from wild type (wt) C57BL/6 and T-cells from Nur77$^{GFP}$ mice on the following days. IL-38 was given (i.p) three times a week. **(A)** Donor-derived immune cells of total CD45+ cells in the liver were determined by flow cytometry (n = 9). **(B)** Profile of donor-derived (H2kb+) immune cells in the liver between the control and IL-38 groups (n = 9). **(C)** Percentages of CD4+ T-cells and CD8+ T-cells of donor-derived T-cells obtained from the liver (n = 9) between the control and IL-38 groups. **(D)** Expression of MHC-II on the surface of the total APCs between the control and IL-38 groups obtained from the liver (n = 9). Each data point corresponds to an individual animal, and the data are from 3 independent experiments. Data are shown as mean ± SEM. *$p$ < 0.05, **$p$ < 0.01, ***$p$ < 0.001, ****$p$ < 0.0001; $p$-values were calculated using multiple $t$-test (B) and unpaired $t$-test (A, C-D).

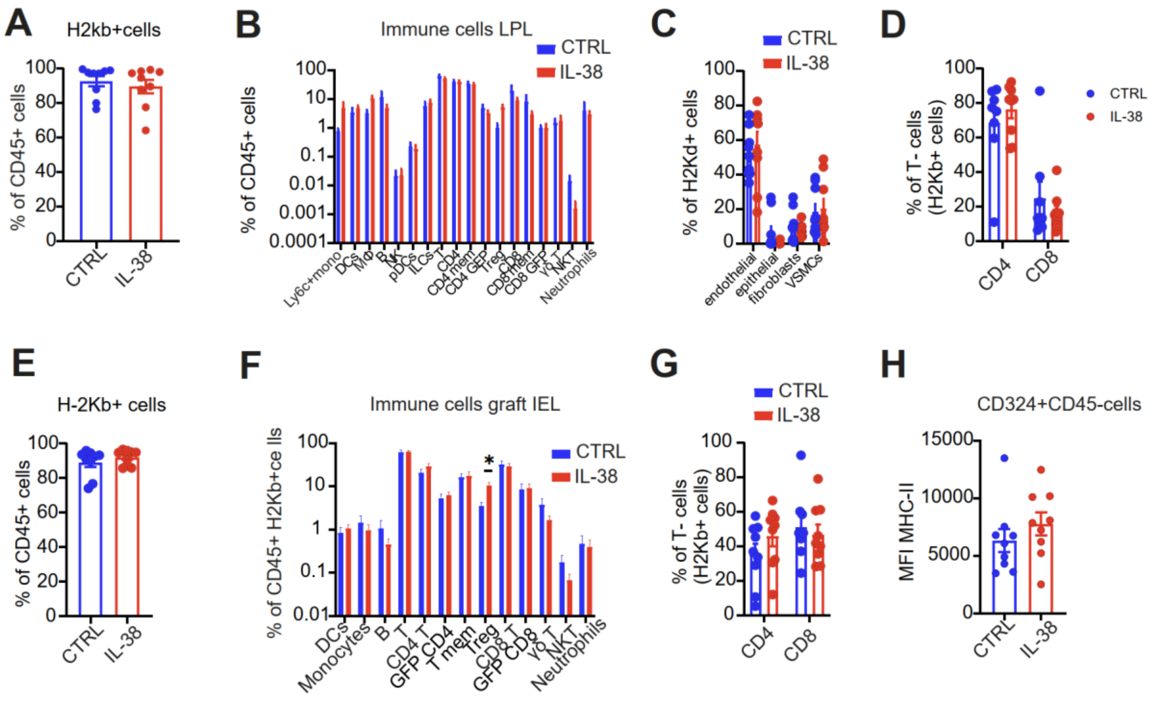


**Figure S15. IL-38 affects immune cells and host-derived cells in the gastrointestinal tract (GI) during GvHD.** BALB/c mice (host) were irradiated on day 0 and received T-cell-depleted bone marrow (BM) cells from wild type (wt) C57BL/6 and T-cells from Nur77$^{GFP}$ mice on the following days. IL-38 was given (i.p) three times a week. **(A)** The proportions of donor-derived immune cells of the total CD45+ cells in the lamina propria (LPL) were determined by flow cytometry (n=9). **(B)**The immunofluorescent profiling of the immune donor-derived (H2kb+) cells in the LPL between the control and IL-38 groups (n=9). **(C)** The percentages of host-derived cells (H2Kd+) presented in the LPL between the control and IL-38 groups (n=9). **(D)** The percentages of CD4 and CD8 T-cells of donor-derived T-cells obtained from the LPL and inspected by flow cytometry (n=9) between the control and IL-38 groups. **(E)** The proportions of donor-derived immune cells of the total CD45+ cells in the intraepithelial lymphocytes (IEL) were determined by flow cytometry (n=9). **(F)**The immunofluorescent profiling of the immune donor-derived (H2kb+) cells in the IEL between the control and IL-38 groups (n=9).**(G)** The percentages of CD4 and CD8 T-cells of donor-derived T-cells obtained from the IEL and inspected by flow cytometry (n=9) between the control and IL-38 groups. **(H)** The expression of MHC-II on the surface of host-derived epithelial cells (CD324+H2Kd+) between control and IL-38 groups obtained in the IEL and inspected by flow cytometry (n=9). Each data point corresponds to an individual animal, and the data are from 3 independent experiments. Data are shown as mean ± SEM. *$p < 0.05$, **$p < 0.01$, ***$p < 0.001$, ****$p < 0.0001$; $p$-values were calculated using multiple $t$-test (B, F) and unpaired $t$-test (A, C, D, E, G, H).

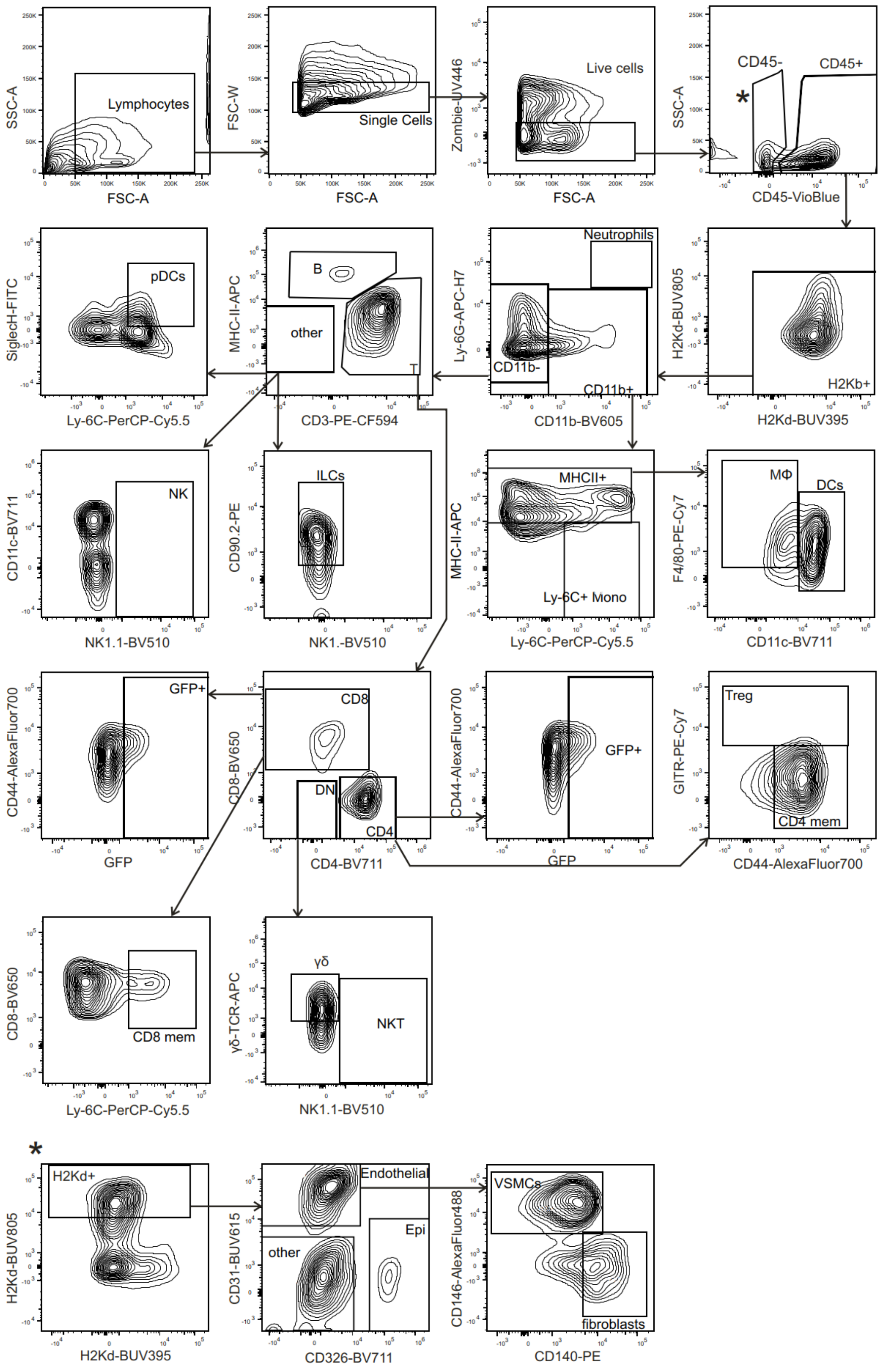


**Figure S16. Representative FACS plots of flow cytometric immunophenotyping of LPL cells from murine GvHD samples.**

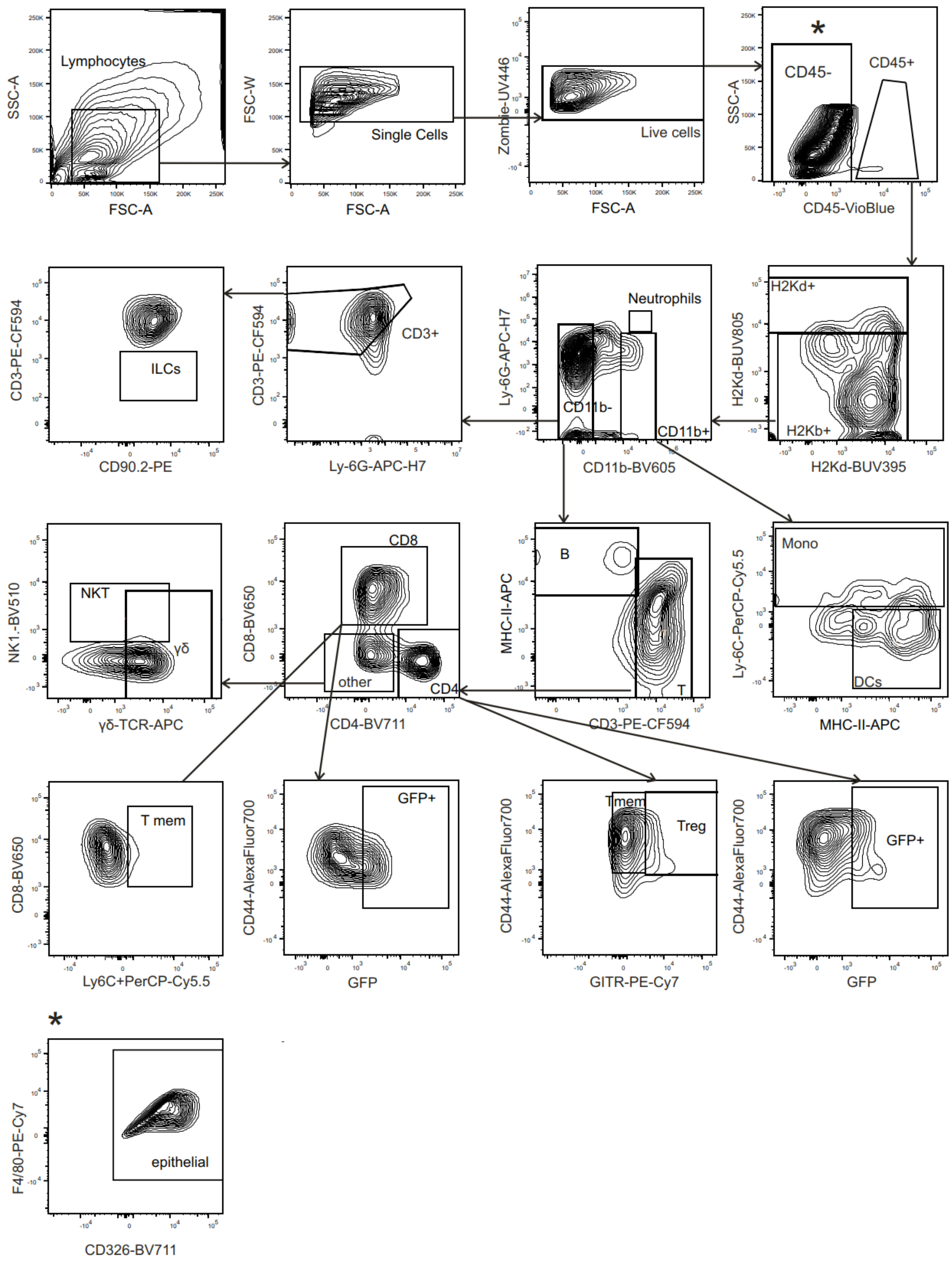


**Figure S17. Representative FACS plots of flow cytometric immunophenotyping of IEL cells from murine GvHD samples.**

### Supplementary materials and methods

#### IL-38 ELISA

IL-38 levels in cell supernatants were quantified using a solid-phase sandwich ELISA (Human IL-38/IL-1F10 DuoSet ELISA, R&D Systems) following the manufacturer's instructions.

#### Reactive Oxygen Species detection

Macrophages were trypsinized and subsequently incubated with CellROX Orange at concentration of 1 µM (Thermo Fisher Scientific) at 37 ºC for 20 min. Afterwards, cells were washed with warm PBS and blocked with human Fc blocking reagent and stained with anti-CD14 Ab. ROS were examined via flow cytometry.

#### Respiration measurement

Respiration in monocytes/macrophages was analyzed using Seahorse XF96 extracellular flux analyzer (Agilent). Cells were plated in Seahorse 96-well cell culture plates at 1 × 10^5 cells/well immediately after isolation, incubated with or without IL-38 for 3 or 7 days, and equilibrated in Seahorse RPMI medium (103576, Agilent) supplemented with 10 mM L-glucose and 2 mM L-glutamine prior to assay. Cells were treated with 2.5 µM oligomycin (Sigma-Aldrich), 1 µM carbonyl cyanide 3-chlorophenylhydrazone (CCCP, Sigma-Aldrich), 1 µg/ml antimycin (Sigma-Aldrich), and 2.5 µM rotenone (Sigma-Aldrich). Oxygen consumption rates (OCR) were monitored in real time after injection of each compound. Maximal respiration rates were defined as OCR after addition of CCCP.

#### Analysis of human GvHD publicly available data sets

Single-cell transcriptomic data sets were downloaded from the Gene Expression Omnibus (GSE191335) and analyzed within Seurat pipeline.
